# Functional disease architectures reveal unique biological role of transposable elements

**DOI:** 10.1101/482281

**Authors:** Farhad Hormozdiari, Bryce van de Geijn, Joseph Nasser, Omer Weissbrod, Steven Gazal, Chelsea J.-T. Ju, Luke O’Connor, Margaux Louise Anna Hujoel, Jesse Engreitz, Fereydoun Hormozdiari, Alkes L. Price

**Affiliations:** Department of Epidemiology, Harvard T.H. Chan School of Public Health, Boston, MA 02115, USA; Program in Medical and Population Genetics, Broad Institute of MIT and Harvard, Cambridge, Massachusetts, USA; Department of Computer Science, University of California, Los Angeles, California 90095; Program in Bioinformatics and Integrative Genomics, Harvard Graduate School of Arts and Sciences, Boston, Massachusetts, USA; Department of Biostatistics, Harvard T.H. Chan School of Public Health, Boston, MA 02115, USA; Department of Biochemistry and Molecular Medicine, University of California, Davis, CA 95616, USA; MIND Institute and UC-Davis Genome Center, Davis, CA 95616, USA

## Abstract

Transposable elements (TE) comprise roughly half of the human genome. Though initially derided as “junk DNA”, they have been widely hypothesized to contribute to the evolution of gene regulation. However, the contribution of TE to the genetic architecture of diseases and complex traits remains unknown. Here, we analyze data from 41 independent diseases and complex traits (average N=320K) to draw three main conclusions. First, TE are uniquely informative for disease heritability. Despite overall depletion for heritability (54% of SNPs, 39±2% of heritability; enrichment of 0.72±0.03; 0.38-1.23 enrichment across four main TE classes), TE explain substantially more heritability than expected based on their depletion for known functional annotations (expected enrichment of 0.35±0.03; 2.11x ratio of true vs. expected enrichment). This implies that TE acquire function in ways that differ from known functional annotations. Second, older TE contribute more to disease heritability, consistent with acquiring biological function; SNPs inside the oldest 20% of TE explain 2.45x more heritability than SNPs inside the youngest 20% of TE. Third, Short Interspersed Nuclear Elements (SINE; one of the four main TE classes) are far more enriched for blood traits (2.05±0.30) than for other traits (0.96±0.09); this difference is far greater than expected based on the weaker depletion of SINEs for regulatory annotations in blood compared to other tissues. Our results elucidate the biological roles that TE play in the genetic architecture of diseases and complex traits.

## Introduction

Transposable elements (TE), defined as DNA sequences that can insert themselves at new genomic locations, comprise roughly half of the human genome ^1,2^. Though initially derided as “junk DNA”, TE have been widely hypothesized to contribute to the evolution of gene regulation by providing new targets for transcription factor binding and rewiring core regulatory networks ^3–16^. TE have been shown to play these important roles in a growing number of specific examples, potentially impacting the genetic architecture of common disease. However, our current understanding of the contribution of TE to the genetic architecture of diseases and complex traits is extremely limited.

Here, we applied stratified LD score regression ^17^ (S-LDSC) with the baseline-LD model ^18^ to 41 independent diseases and complex traits (average N=320K) to estimate the components of heritability explained by different classes of TE. We sought to answer three questions. First, what is the contribution of TE to disease, and does this differ from what is expected based on the extent of their level of overlap with known functional annotations ^19,20^? Second, do older TE contribute more to the disease heritability than younger TE? Third, do there exist classes of TE that play a greater role in specific diseases or traits?

## Results

### Overview of methods

We applied stratified LD score regression (S-LDSC) ^17^ to assess the contribution of different TE to disease and trait heritability. We estimated the heritability enrichment and standardized effect size (*τ*\*) for each TE annotation conditional on 75 functional annotations from the baseline-LD model ^18^ (Supplementary Table 1, see URLs). Heritability enrichment is defined as the proportion of heritability causally explained by the set of common SNPs in an annotation divided by the proportion of common SNPs in the annotation. Distinct from heritability enrichment, we also compute the enrichment that is expected based on an annotation’s overlap with baseline-LD model annotations, denoted as Expected (baseline-LD) (see Methods); this computation determines the extent to which heritability enrichment/depletion is explained by known functional annotations. We note that enrichment and expected enrichment can be either > 1 or < 1 (i.e. depletion). Standardized effect size (*τ*\*) is defined as the proportionate change in per-SNP heritability associated with an increase in the value of the annotation by one standard deviation ^18^; unlike heritability enrichment, *τ*\* quantifies effects that are unique to the focal annotation (see Methods). For each TE annotation, we include an additional annotation defined by 500bp flanking regions, to guard against bias due to model misspecification ^17^ (see Methods). We have made our annotations and partitioned LD scores freely available (see URLs). Most of our results are meta-analyzed across 41 independent diseases and complex traits (Supplementary Table 2, same traits as in ref.^21^).

### TE are uniquely informative for disease heritability

We first focused on four main TE classes: long interspersed nuclear elements (LINE; 21% of SNPs), short interspersed nuclear elements (SINE; 16% of SNPs), long terminal repeats (LTR; 9.8% of SNPs), DNA transposons (DNA; 3.2% of SNPs), and the union all TE (ALLTE; 54% of SNPs). The proportion of SNPs in each TE class slightly exceeded the proportion of the genome spanned by the TE class (Supplementary Figure 1). This is consistent with weaker selective constraint within surviving TE, and confirms that SNPs lying inside TE can be effectively assayed despite the challenges of aligning TE sequences. ALLTE explained 39% of disease heritability (meta-analyzed across 41 diseases and traits), a moderate depletion (enrichment of 0.72±0.03; Figure 1A-B and Supplementary Table 3). The four main TE classes were all depleted or non-significantly enriched for trait heritability, with substantial heterogeneity between classes: 0.73±0.05 for LINE, 1.18±0.11 for SINE, 0.38±0.07 for LTR, and 1.23±0.19 for DNA (Figure 1B and Supplementary Table 3). Our simulations confirm that S-LDSC produces unbiased estimates of enrichment for these annotations (see Methods, Supplementary Figure 2). A secondary analysis of enrichment of fine-mapped causal disease SNPs ^22,23^ produced concordant results (Supplementary Table 4). A secondary analysis of enrichment of fine-mapped causal cis-eQTL SNPs ^21^ from GTEx data ^24^ also produced concordant results (Supplementary Table 5).

**Figure 1.**
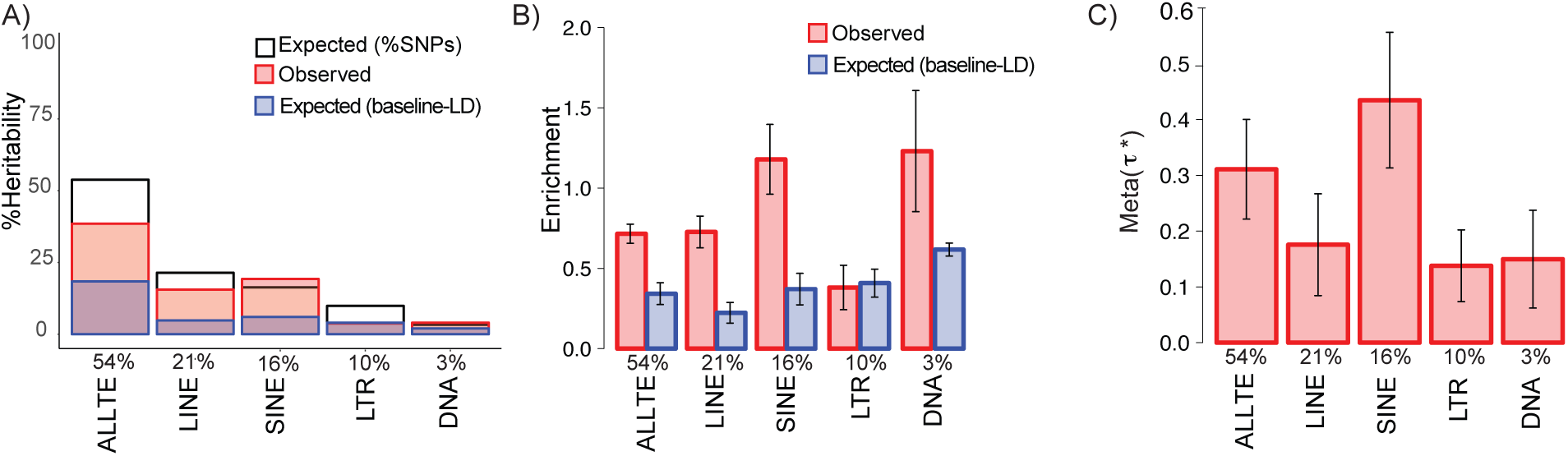
TE are uniquely informative for disease heritability. For each of four main TE classes and ALLTE, we report A) three measures of %heritability: Expected (%SNPs), Observed, and Expected (baseline-LD); B) two measures of heritability enrichment: Observed and Expected (baseline-LD); and *τ*\* standardized effect size (*τ*\*), which quantifies effect that are unique to the focal annotation. Results are meta-analyzed across 41 independent traits. Numerical values of %SNPs are provided for each annotation. Error bars denote 95% confidence intervals. Numerical results are reported in Supplementary Table 3.

Notably, the heritability enrichments expected based on overlap with baseline-LD model annotations were much lower (Expected (baseline-LD); Figure 1A-B and Supplementary Table 3), consistent with the large depletion of overlap between TE and known functional annotations (Supplementary Figure 3). Accordingly, *τ*\*estimates were significantly positive for each TE class (Figure 1C), implying disease heritability enrichment effects that are not captured by known functional annotations. These *τ* estimates were similar (in absolute value) to *τ* estimates for the most informative annotations in our previous work ^18,21^. Furthermore, each of the four main TE classes (LINE, SINE, LTR, and DNA) had a significant *τ* conditional on each other and baseline-LD model annotations (Supplementary Table 6), indicating that each is uniquely informative for disease heritability.

We investigated whether the age of a TE impacts its contribution to disease heritability. We estimated the age of each TE using miliDiv (RepeatMasker software; see URLs), which computes the number of mutations relative to a consensus sequence to estimate the age of each TE^11^. We stratified SNPs lying in a TE into five quintiles based on the age of the TE. We determined that older SNPs had larger heritability enrichments than younger SNPs (e.g. 0.91±0.11 for oldest quintile vs 0.37±0.10 for youngest quintile; Supplementary Figure 4 and Supplementary Table 7). We repeated this analysis for each TE class (SINE, LINE, LTR, DNA) and observed the largest effect for SINE (Supplementary Figure 5). Analyses of Expected(baseline-LD) across TE families/subfamilies produced similar results, with the largest age effect for SINE (Supplementary Figure 6 and Supplementary Table 8). These results indicate that older TE have a higher contribution to disease heritability, perhaps because they have gained biological function.

Next, we analyzed 35 TE families/subfamilies spanning at least 0.4% of common SNPs (i.e., MAF 0.05; Supplementary Table 9). We identified 4 TE families/subfamilies that were significantly depleted for trait heritability (L1, L1PA3, ERV1, and L1PA4; Supplementary Figure 7 and Supplementary Tables 10 and 11); none were significantly enriched. For the 814 TE families/subfamilies spanning less than 0.4% of common SNPs (Supplementary Table 12), we estimated Expected (baseline-LD) enrichment only (Supplementary Table 13), as S-LDSC is not applicable to very small annotations ^17^. We identified 587 TE families/subfamilies that were significantly depleted for expected disease heritability (Supplementary Table 14). We also identified 46 TE families/subfamilies that were significantly enriched for expected disease heritability (Supplementary Figure 8 and Supplementary Table 14), consistent with their excess overlap with known functional annotations (Supplementary Figures 9 and 10 and Supplementary Tables 15 and 16). Notably, LFSINE-Vert and AmnSINE1, which have previously been reported to have important biological function ^25–27^, had very large expected enrichments (5.54±0.39 and 5.44±0.32 respectively).

### SINE are specifically strongly enriched for blood traits

We investigated whether TE enrichment varies across disease and traits. We estimated the heritability enrichment of each TE class (ALLTE, LINE, SINE, LTR, DNA) for 5 blood traits, 6 autoimmune diseases, and 8 brain-related traits (see Supplementary Table 17; same traits as in ref. ^21^). We included a blood-specific chromatin annotation in our analyses of blood traits and autoimmune diseases, and a brain-specific chromatin annotation in our analyses of brain-related traits, in addition to the baseline-LD model (see Methods). Results are reported in Figure 2A and Supplementary Table 18. We determined that SINE are specifically strongly enriched for blood traits (2.05±0.30 vs. 1.18± 0.11 for non-blood traits; P=3E-04 for difference); no other TE class had significant trait class-specific enrichment after correcting for hypotheses tested, although SINE enrichment was non-significantly higher for autoimmune diseases vs. other traits (Supplementary Table 18). The difference in SINE enrichment for blood traits vs non-blood traits was much higher than expected based on overlap with baseline-LD model and blood-specific chromatin annotations (Expected (baselineLD+blood chromatin); Supplementary Table 18). Accordingly, we estimated a particularly large *τ* for SINE for blood traits (Figure 2B and Supplementary Table 18), much larger (in absolute value) than *τ* estimates for the most informative annotations in our previous work ^18,21^. The specific importance of SINE for blood traits is consistent with the weaker depletion of SINE in blood-specific chromatin annotations vs. other tissue/cell types (Figure 2C and Supplementary Table 19), but is far greater than expected based on this weaker depletion; in particular, the *τ* estimates of Figure 2B are conditioned on blood-specific chromatin annotations.

**Figure 2.**
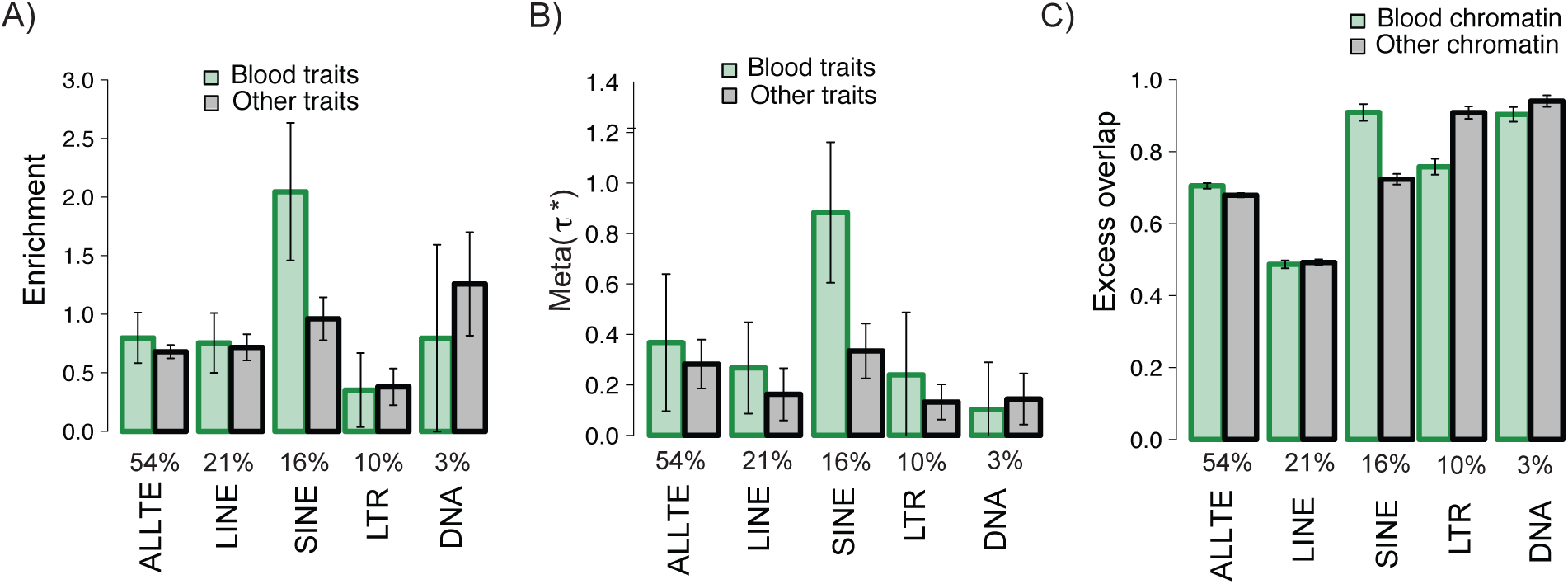
Larger SINE enrichments for blood traits. For each of four main TE classes and ALLTE, we report A) heritability enrichment for blood traits and other traits; B) standardized effect size (*τ*\*) for blood traits and other traits; and C) excess overlap with chromatin annotations in blood and chromatin annotations in other tissues. Results are meta-analyzed across 41 independent traits. Numerical values of %SNPs are provided for each annotation. Error bars denote 95% confidence intervals. Numerical results are reported in Supplementary Table 18 (A,B) and Supplementary Table 19 (C).

We repeated the trait class-specific analysis for the 35 TE families/subfamilies spanning at least 0.4% of common SNPs (Supplementary Tables 20-22). We did not detect any trait class-specific enrichments except for the Alu family, which spans 80% of the SINE class and produces results similar to SINE. For the 814 TE families/subfamilies spanning less than 0.4% of common SNPs, we detected 27 that had significantly higher Expected (baseline-LD+blood chromatin) enrichment for blood-related traits vs. other traits (Supplementary Table 23) and 27 that had significantly higher Expected (baseline-LD+blood chromatin) enrichment for autoimmune diseases vs other traits (Supplementary Table 24). The majority of TE families/subfamilies for that were specifically enriched for autoimmune diseases are endogenous retroviruses (ERV), including the MER41, which has previously been reported to contribute to autoimmune disease ^14^. We also detected 109 TE families/subfamilies with higher Expected (baselineLD+brain chromatin) enrichment for brain-related traits vs. other traits (Supplementary Table 25).

## Discussion

We have quantified the disease heritability explained by TE, including different classes of TE. We reached three main conclusions. First, TE are uniquely informative for disease heritability, as they explain substantially more heritability than expected based on their depletion for known functional annotations. This implies that TE acquire function in ways that differ from known functional annotations. Second, we observed that older TE contribute more to disease heritability, consistent with acquiring biological function. Third, the SINE class of TE is far more enriched for blood traits than for other traits, showing that TE biology can be trait class-specific.

Our findings have several biological implications. First, our results suggest that the functional annotation of the human genome is far from complete, as the functional regions underlying the contribution of TE to disease heritability have yet to be annotated. This motivates intense efforts to identify these functional regions. We have provided a framework (*τ*\*metric; Figure 1C) to evaluate these efforts. Specifically, a *τ\** value close to 0 (conditional on a new set of functional annotations) would imply that this goal has been achieved; this can be evaluated for all TE and all traits (Figure 1C), but is of particular interest for SINE and blood traits (Figure 2B). Second, our TE-related annotations with conditionally significant signals (Figure 1C) can be incorporated to improve functionally informed fine-mapping ^28–30^, as well as functionally informed efforts to increase association power ^31,32^ and polygenic prediction accuracy^33–35^.

We note several limitations of our work. First, S-LDSC cannot be applied to estimate the heritability enrichment of TE families/subfamilies that span a small proportion of the genome (e.g. less than 0.4% of common SNPs) ^17^. We can instead compute the heritability enrichment that is expected based on an annotation’s overlap with baseline-LD model annotations, although we caution that this quantity has a different interpretation. Second, we focused our analyses on common variants, as we used the 1000 Genomes LD reference panel, but future work could draw inferences about low-frequency variants using larger reference panels ^36^. Third, SNPs lying inside TE may be difficult to identify and annotate due to the challenges of aligning TE sequences. However, the proportion of SNPs in each TE class slightly exceeded the proportion of the genome spanned by the TE class (Supplementary Figure 1), suggesting that SNPs lying inside TE can be effectively assayed.

In addition, we observed that 85% of 1000 Genomes SNPs lie in a 35-mer that has mappability of 1 (i.e. unique mappability) based on the ENCODE 35-mer track. Thus, 85% of the SNPs in our analysis are not impacted by aligning reads to repetitive regions of the genome. Furthermore, we confirmed that restricting our analyses to the 85% of SNPs with mappability of 1 produces very similar results (Supplementary Table 27). Fourth, inferences about components of heritability can potentially be biased by failure to account for LD-dependent architectures ^18,37–39^. All of our analyses used the baseline-LD model, which includes 6 LD-related annotations ^18^. The baseline-LD model is supported by formal model comparisons using likelihood and polygenic prediction methods, as well as analyses using a combined model incorporating alternative approaches ^40^; however, there can be no guarantee that the baseline-LD model perfectly captures LD-dependent architectures. Despite these limitations, our results substantially improve our current understanding of the contribution of TE to the genetic architecture of diseases and complex traits.

## Supporting information

## Acknowledgements

We are grateful to Armin Schoech and Po-Ru Loh for helpful discussions. This research was funded by NIH grants U01 HG009379, R01 MH101244, R01 MH109978 and R01 MH107649. This research was conducted using the UK Biobank Resource under Application 16549. F.H. is also supported by NIH grants T32 DK110919 and F32HG009987.

## Methods

### Heritability enrichment and standardized effect size (*τ*\*)

We use two metics (Heritability enrichment and standardized effect size (*τ\**)) to measure the contribution of an annotation to disease and trait heritability ^17,18^. We use S-LDSC to compute the heritability enrichment and standardized effect size (*τ\**). S-LDSC assumes that the per-SNP heritability or variance of each SNP is equal to the linear contribution of each annotation ^17^:

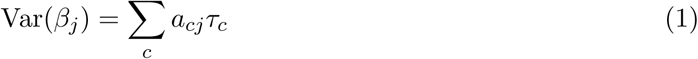

where *a*_*cj*_ indicates the annotation value of SNP *j* for the annotation *c* and *τ*_*c*_ is the contribution of annotation *c* to the per-SNP heritability. S-LDSC estimates the *τ*_*c*_ for each annotation using the following equation:

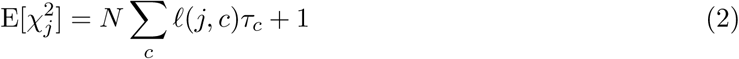

where *N* is GWAS sample size and *ℓ* (*j, c*) is the LD-score for the SNP *j* and annotation c computed from the 1000 Genome project (see URLs). We estimated *ℓ*(*j, c*) as 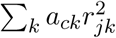 where *r*_*jk*_ is the genotypic correlation between SNPs *j* and *k*.

Because *τ*_*c*_ depends on trait heritability and the size of annotation we can not compare *τ*_*c*_ between different traits or annotations. Gazal et al.^18^ introduced standardized effect size (*τ\**) for an annotation as follows:

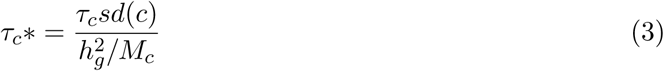

where *sd*(*c*) is the standard deviation of the annotation values, *M*_*c*_ is total number of common SNPs used to estimate the 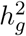, and 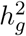 is the SNP-heritability for each trait. In our experiments *M*_*c*_ is equal to 5,961,159. We can compare *τ\** between different traits or annotations.

Heritability enrichment for an annotation is defined as the proportion of trait heritability captured by an annotation divide by the proportion of common SNPs that span that annotation. Thus, heritability enrichment is computed as follows:

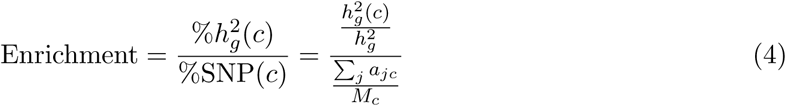

where 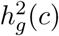 is the heritability captured by the annotation *c* and it is computed as follows:

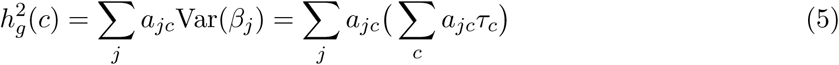

Both heritability enrichment and *τ\** are computed conditional on set of annotations in the model (e.g. baseline-LD model ^18^), *τ\** captures the signal that is unique to the focal annotation after conditioning on all the annotations in the model. Standardized effect size (*τ\**) is defined as the proportionate change in per-SNP heritability associated with an increase in the value of the annotation by one standard deviation ^18^. However, enrichment captures a signal that is unique and/or non-unique to the focal annotation.

We computed the statistical significance of heritability enrichment using block-jackknife, as described in our previous studies ^17,18,21^ where we break the genome to 200 equal blocks. We compute the statistical significant of *τ\** by assuming that 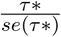 follows a normal distribution with mean zero and variance one 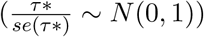^17,18,21^

The meta-analyzed values of enrichment and *τ\** across the 41 independent traits (47 data sets, see Supplementary Table 2) were computed using a random-effect meta-analysis, as implemented in the rmeta R package (see URLs).

#### Expected(baseline-LD)

We compute the expected (baseline-LD) enrichment for an annotation by assuming that the *τ* of the focal annotation is zero. This is equivalent to apply S-LDSC to each trait using baseline-LD model and compute the per-SNP heritability for each variant using equation (1). Next, we compute the 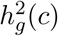 of the annotation by summing over the per-SNP heritability of all SNPs that are in the annotation c. In the end, we can compute the heritability enrichment using equation (4) and compute the standard error using similar block-jackknife.

### Proportion of heritability captured by each annotation

We can compute 3 different proportion of heritability (%heritability) for each annotation: observed heritability, Expected(%SNPs), Expected (baseline-LD). Observed heritability enrichment is obtained from S-LDSC by conditioning on baseline-LD model.

#### Expected(%SNPs)

Under the null model, we assume the enrichment of an annotation is one. Thus, this annotation has none significant heritability enrichment (non-enriched or non-depleted), then we expect the %heritability for an annotation to be equal to %SNPs in that annotation.

#### Expected(baseline-LD)

We compute this quantity by multiplying the expected (baseline-LD) and%SNPs.

#### Observed %heritability

We compute this quantity by utilizing S-LDSC results of heritability enrichment and %SNPs. We have: Observed %heritability = Heritability enrichment × %SNPs.

### TE annotations

We constructed two annotations for each TE where the first annotation is obtained by considering all the SNPs that fall in a TE and the second annotation is obtained by considering all the SNP in a 500bp window of the TE. The window annotation is based on recommendation of previous work ^17^. All results are obtained by conditioning over baseline-LD model. The *τ\** and enrichment reported for each TE class/family/subfamily are based on the first constructed TE annotation. We compared this enrichment estimates with the case where we compute the enrichment of an annotation conditional jointly on 4 extra annotations created by considering different window size of 100, 200, 500, and 1000bp. We observed that S-LDSC results does not depend on the window size (Supplementary Figure 11).

### S-L DSC simulations

We set the *τ* for each annotation based on enrichment obtained in real data sets. Utilizing the total heritability we simulated causal trait effect sizes using a polygenetic model: 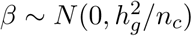 where *n*_*c*_ is the number of causal SNPs. We simulated the phenotypic values under the additive model (*Y* = *X*(*β* + *e*) where X is the standardized genotype matrix and *e* is the environment and measurement noise. We computed the summary statistics by performing linear regression between the phenotypic values and genotype data using PLINK software (see URLs). In our simulation, we vary the number of individuals for the traits among 2,000, 20,000, and 40,000 where UK biobank genotypes ^41^ are used. After simulating the summary statistics, we applied S-LDSC conditional on baseline-LD model and our TE annotation. Regression SNPs in S-LDSC were obtained from the HapMap Project phase 3^42^ (see URLs). These SNPs are well-imputed SNPs. SNPs with marginal association statistics larger than 80 or larger than 0.001N and SNPs that are in the major histocompatibility complex (MHC) region were excluded from all the analyses ^17,18,21^. Reference SNPs were obtained using the European samples in 1000G ^41^. Heritability SNPs, which are used to estimate 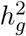, were common variants (MAF≥0.05) in the set of reference SNPs.

### Excess overlap

Let *A* and *B* indicate two annotations and |.| indicate the number of SNPs in the annotation. We defined the excess overlap as follows:

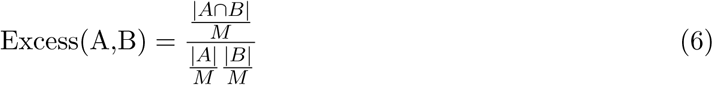

where *M* is total number of SNPs and |*A* ⋂ *B*| indicates the set of SNPs that is shared in both annotations *A* and *B*. We compute the standard error over our estimates using block jackknife with 200 blocks that is similar how S-LDSC computed the standard error over heritability enrichment as described in our previous studies ^17,18,21^.

### Tissue-specific chromatin annotations

Blood chromatin is blood active chromatin regions by combining 27 blood cells and 6 chromatin marks (H3K27ac, H3K4me3, DNase, DNase-H3K27ac, DNase-H3K4me3) obtained from ChromImpute ^43^ applied on Roadmap Epigenomics data ^20^. Non-blood chromatin is non-blood active chromatin regions by combining 100 non-blood cells and 6 chromatin marks (H3K27ac, H3K4me3, DNase, DNase-H3K27ac, DNase-H3K4me3).

Brain chromatin is brain active chromatin regions by combining 13 brain cells and 6 chromatin marks (H3K27ac, H3K4me3, DNase, DNase-H3K27ac, DNase-H3K4me3) obtained from ChromImpute ^43^ applied on Roadmap Epigenomics data ^20^. Non-brain chromatin is non-brain active chromatin regions by combining 114 non-brain cells and 6 chromatin marks.

### URLs

baselineLD annotations https://data.broadinstitute.org/alkesgroup/LDSCORE/

TE annotations: https://data.broadinstitute.org/alkesgroup/LDSCORE/TE/

Repeat masker software: http://www.repeatmasker.org

1000 Genomes Project Phase 3 data: ftp://ftp.1000genomes.ebi.ac.uk/vol1/ftp/release/20130502

PLINK software: https://www.cog-genomics.org/plink2

BOLT-LMM software: https://data.broadinstitute.org/alkesgroup/BOLT-LMM

BOLT-LMM summary statistics for UK Biobank traits: https://data.broadinstitute.org/alkesgroup/UK

UK Biobank: http://www.ukbiobank.ac.uk/

UK Biobank Genotyping and QC Documentation: http://www.ukbiobank.ac.uk/wp-content/uploads/201404/UKBiobank_genotyping_QC_documentation-web.pdf

rmeta R package: https://cran.r-project.org/web/packages/rmeta/index.html

