## Supplementary material for "Functional disease architectures reveal unique biological role of transposable elements"

### Supplementary Tables

| Annotation | #Source | Model |
|---|---|---|
| base | Finucane et al.[17] | baseline/baselineLD |
| Coding | UCSC Genome Browser and Finucane et al.[17] | baseline/baselineLD |
| 3' UTR | UCSC Genome Browser and Finucane et al.[17] | baseline/baselineLD |
| 5' UTR | UCSC Genome Browser and Finucane et al.[17] | baseline/baselineLD |
| Promoter | UCSC Genome Browser and Finucane et al.[17] | baseline/baselineLD |
| Intron | UCSC Genome Browser and Finucane et al.[17] | baseline/baselineLD |
| ENCODE TF binding | ENCODE[44] and Finucane et al.[17] | baseline/baselineLD |
| CTCF promoter | Hoffman et al[45] and Finucane et al.[17] | baseline/baselineLD |
| Promotor-Flanking | Hoffman et al[45] and Finucane et al.[17] | baseline/baselineLD |
| Transcribed | Hoffman et al[45] and Finucane et al.[17] | baseline/baselineLD |
| Transcription start size (TSS) | Hoffman et al[45] and Finucane et al.[17] | baseline/baselineLD |
| Strong enhancer | Hoffman et al[45] and Finucane et al.[17] | baseline/baselineLD |
| Weak enhancer | Hoffman et al[45] and Finucane et al.[17] | baseline/baselineLD |
| DHS | ENCODE[44]/Roadmap[20] post-processed by Trynka et al and Finucane et al.[17] | baseline/baselineLD |
| H3K4me1 | Roadmap Epigenomics[20] post-processed by Trynka et al[28] and Finucane et al.[17] | baseline/baselineLD |
| H3K4me3 | Roadmap Epigenomics[20] post-processed by Trynka et al[28] and Finucane et al.[17] | baseline/baselineLD |
| H3K9ac | Roadmap Epigenomics[20] post-processed by Trynka et al[28] and Finucane et al.[17] | baseline/baselineLD |
| H3K27ac | Roadmap Epigenomics[20] post-processed by Trynka et al[28] and Finucane et al.[17] | baseline/baselineLD |
| H3K27ac | Hnisz et al[46] and Finucane et al.[17] | baseline/baselineLD |
| Super Enhancer | Hnisz et al[46] and Finucane et al.[17] | baseline/baselineLD |
| Conserved | Lindblad-Toh et al.[47] and Finucane et al.[17] | baseline/baselineLD |
| FANTOM5 enhancer | Anderson et al.[48] and Finucane et al.[17] | baseline/baselineLD |
| 2 GERP[49] conserved | Gazal et al.[18] | baselineLD |
| 10 MAF bin | Gazal et al.[18] | baselineLD |
| Predicted Allele Age | Gazal et al.[18] | baselineLD |
| LLD-AFR | Gazal et al.[18] | baselineLD |
| Recombination Rate | Gazal et al.[18] | baselineLD |
| Nucleotide Diversity | Gazal et al.[18] | baselineLD |
| Background Selection Statistics | Gazal et al.[18] | baselineLD |
| CpG-Content | Gazal et al.[18] | baselineLD |

**Supplementary Table 1. List of 75 functional and LD related annotations in the baselineLD model (v1.1).** There are 24 annotations in the baselineLD where we add a 500bp window around the category. In the case of DHS, H3K4me1, H3K4me3, and H3K27ac annotations we add 100-bp window.

See attached Excel file

**Supplementary Table 2. List of 47 data sets analyzed in this work.** We obtained publicly available summary statistics from previous published studies. UK Biobank summary statistics were previously computed using BOLT-LMM[50,51]. For some traits we have more than one data set, thus we have 41 independent traits. However, we utilized all 47 data sets in our meta-analyses as the number of samples that overlap is low. These traits were selected based on a heritability z-score > 6. The 41 traits and 47 data sets are identical to those analyzed in ref.[21].

| TE | Expected (%SNPs) | $\tau*(se)$ | $\tau*$ (P) | Enrichment (se) | Enrichment (P) | Observed | Expected (baseline-LD) |
|---|---|---|---|---|---|---|---|
| ALLTE | 53.81 | 0.31 (0.05) | 1.14E-11 | 0.72 (0.03) | 3.68E-12 | 38.52 | 18.42 |
| LINE | 21.41 | 0.18 (0.05) | 1.73E-04 | 0.73 (0.05) | 4.66E-07 | 15.57 | 4.78 |
| SINE | 16.32 | 0.43 (0.06) | 3.09E-12 | 1.18 (0.11) | 2.73E-01 | 19.25 | 6.05 |
| DNA | 3.28 | 0.15 (0.04) | 8.64E-04 | 1.23 (0.19) | 6.68E-01 | 4.04 | 4.03 |
| LTR | 9.88 | 0.14 (0.03) | 2.90E-05 | 0.38 (0.07) | 2.81E-08 | 3.76 | 2.02 |

**Supplementary Table 3. S-LDSC results for four main TE classes and ALLTE.** We compute the heritability enrichment and $\tau*$ for four main TE classes (LINE, SINE, LTR, and DNA) and ALLTE (combination of all TE) conditional on baseline-LD model. These results are obtained by meta-analyzing 41 independent traits and disease.

| TE | Enrichment (Farh et al.) | Enrichment (Huang et al.) | S-LDSC |
|---|---|---|---|
| ALLTE | 0.81 (0.01) | 0.92 (0.03) | 0.72(0.03) |
| LINE | 0.74 (0.03) | 0.79 (0.08) | 0.73(0.05) |
| SINE | 1.08 (0.04) | 1.23 (0.13) | 1.18(0.11) |
| LTR | 0.64 (0.03) | 0.77 (0.11) | 0.38(0.07) |
| DNA | 1.01 (0.09) | 1.07 (0.16) | 1.23(0.19) |

**Supplementary Table 4. Enrichment of fine-mapped causal disease SNPs for four main TE classes and ALLTE.** We computed the enrichment of fine-mapped causal disease SNPs as the proportion of fine-mapped SNPs (from Farh et al.[22] and Huang et al.[23]) that line in TE divide by the proportion of common variants that lie in TE. This is a conservative computation (results may be biased towards 1) because fine-mapped SNPs were defined based on 95% causal sets and may include SNPs that are not actually causal. We also report the disease heritability enrichment estimated using S-LDSC.

| TE | Enrichment (fine-mapping) FE-Meta-Tissue | Enrichment (fine-mapping) Whole Blood | S-LDSC |
|---|---|---|---|
| ALLTE | 0.89 (0.01) | 0.88 (0.01) | 0.72(0.03) |
| LINE | 0.67 (0.01) | 0.61 (0.01) | 0.73(0.05) |
| SINE | 1.30 (0.02) | 1.34 (0.02) | 1.18(0.11) |
| LTR | 0.78 (0.01) | 0.78 (0.01) | 0.38(0.07) |
| DNA | 0.87 (0.02) | 0.82 (0.03) | 1.23(0.19) |

**Supplementary Table 5. Enrichment of fine-mapped causal cis-eQTL SNPs for four main TE classes and ALLTE.** We computed the enrichment of fine-mapped causal cis-eQTL SNPs as the average MaxCPP (maximum Causal Posterior Probability) score[21] of variants that lie in TE divided by the average MaxCPP score of all common SNPs. We obtained MaxCPP scores for FE-Meta-Tissue and Whole blood from ref.[21]. We also report the disease heritability enrichment estimated using S-LDSC.

| TE | Expected (%SNPs) | $\tau*(se)$ | $\tau*$ (P) | Enrichment (se) | Enrichment (P) |
|---|---|---|---|---|---|
| LINE | 21.41 | 0.25 (0.04) | 2.19E-08 | 0.67 (0.04) | 3.63E-08 |
| SINE | 16.32 | 0.48 (0.07) | 4.57E-12 | 1.13 (0.11) | 4.67E-01 |
| DNA | 3.28 | 0.20 (0.04) | 2.21E-07 | 1.20 (0.18) | 5.93E-01 |
| LTR | 9.88 | 0.21 (0.03) | 8.84E-10 | 0.35 (0.07) | 8.80E-08 |

**Supplementary Table 6. S-LDSC results for joint analysis of four main TE classes.** We compute the heritability enrichment and $\tau*$ for four main TE classes (LINE, SINE, LTR, and DNA) conditional on each other and baseline-LD model. These results are obtained by meta-analyzing 41 independent traits and disease.

| Quintile | Expected (baseline-LD) | Observed Enrichment (se) | Observed $\tau*(se)$ |
|---|---|---|---|
| Q1 | 0.01 (0.04) | 0.37 (0.10) | -0.02 (0.03) |
| Q2 | 0.17 (0.05) | 0.63 (0.09) | 0.05 (0.03) |
| Q3 | 0.24 (0.04) | 0.65 (0.06) | 0.01 (0.03) |
| Q4 | 0.50 (0.02) | 0.99 (0.08) | 0.05 (0.03) |
| Q5 | 0.77 (0.02) | 0.91 (0.11) | -0.10 (0.04) |

**Supplementary Table 7. S-LDSC results for all TE in different age quintiles.** We report the Expected (baseline-LD) enrichment and Observed enrichment and $\tau*$ of SNPs in ALLTE for five different age quintiles where Q1 is the youngest and Q5 is the oldest. We computed the Observed enrichment and $\tau*$ conditional on ALLTE and the baseline-LD model. Error bars represent 95% confidence intervals.

See attached Excel file

**Supplementary Table 8. S-LDSC results for 854 TE classes/families/subfamilies.** We computed the Expected (baseline-LD) 853 TE that capture at least 0.001% of common SNPs. These results are obtained by meta-analyzing 41 independent traits and disease. We fit the baselineLD model for each trait. Then, using S-LDSC model we can estimate the per SNP heritability (variance of each SNP). We compute the heritability of each TE by summing over the per SNP heritability of all SNPs in each corresponding TE. The miliDiv column is obtained from Repeat masker software (see URLs). We use the miliDiv as an approximation of the age of the repeat.

| **TE** | **%SNP** |
|---|---|
| ALU | 13.59% |
| AluJb | 1.19% |
| AluJo | 0.58% |
| AluJr | 0.65% |
| AluSc | 0.47% |
| AluSg | 0.56% |
| AluSp | 0.66% |
| AluSq2 | 0.70% |
| AluSx | 1.53% |
| AluSx1 | 1.37% |
| AluSz | 1.15% |
| AluSz6 | 0.47% |
| AluY | 2.22% |
| ERV1 | 3.35% |
| ERVL | 5.98% |
| ERVL-MaLR | 3.96% |
| hAT-Charlie | 1.50% |
| L1 | 17.56% |
| L1M5 | 0.53% |
| L1ME1 | 0.49% |
| L1PA3 | 0.93% |
| L1PA4 | 0.94% |
| L1PA5 | 0.75% |
| L1PA6 | 0.45% |
| L1PA7 | 0.71% |
| L1PB1 | 0.44% |
| L2 | 0.49% |
| L2 | 3.41% |
| L2a | 1.42% |
| L2b | 0.62% |
| L2c | 0.88% |
| MIR | 0.89% |
| MIRb | 1.10% |
| MIRc | 0.41% |
| TcMar-Tigger | 1.13% |

**Supplementary Table 9. List of all 35 TE families/subfamilies spanning at least 0.4% of common SNPs.**

| TE | %SNP | $\tau*(se)$ | $\tau*$ (P) | Enrichment (se) | Enrichment (P) |
|---|---|---|---|---|---|
| ALU | 13.59% | 0.49(0.06) | 1.12E-15 | 1.18(0.12) | 2.35E-01 |
| AluJb | 1.19% | -0.01(0.05) | 8.39E-01 | 0.79(0.35) | 1.26E-01 |
| AluJo | 0.58% | 0.02(0.05) | 6.12E-01 | 1.38(0.52) | 5.41E-01 |
| AluJr | 0.65% | 0.00(0.05) | 9.67E-01 | 1.14(0.62) | 9.53E-01 |
| AluSc | 0.47% | 0.02(0.05) | 7.23E-01 | 0.63(0.54) | 9.75E-03 |
| AluSg | 0.56% | -0.06(0.05) | 2.05E-01 | -0.36(0.56) | 1.65E-01 |
| AluSp | 0.66% | 0.13(0.06) | 3.02E-02 | 1.76(0.60) | 1.36E-01 |
| AluSq2 | 0.70% | -0.01(0.06) | 8.02E-01 | 0.59(0.56) | 9.20E-01 |
| AluSx | 1.53% | 0.05(0.05) | 3.23E-01 | 0.92(0.37) | 9.11E-01 |
| AluSx1 | 1.37% | 0.06(0.05) | 1.84E-01 | 1.00(0.30) | 2.20E-01 |
| AluSz | 1.15% | 0.04(0.05) | 3.35E-01 | 1.01(0.34) | 9.20E-01 |
| AluSz6 | 0.47% | 0.02(0.05) | 7.14E-01 | 0.63(0.54) | 8.03E-02 |
| AluY | 2.22% | -0.03(0.06) | 6.45E-01 | 0.18(0.29) | 5.68E-03 |
| ERV1 | 3.35% | 0.16(0.03) | 2.11E-06 | 0.37(0.09) | 9.08E-07 |
| ERVL | 5.98% | 0.16(0.04) | 2.09E-04 | 0.62(0.11) | 2.82E-02 |
| ERVL-MaLR | 3.96% | 0.09(0.04) | 3.39E-02 | 0.43(0.16) | 9.39E-03 |
| hAT-Charlie | 1.50% | 0.10(0.04) | 1.66E-02 | 1.21(0.33) | 9.66E-01 |
| L1 | 17.56% | 0.16(0.04) | 3.23E-04 | 0.62(0.05) | 2.18E-10 |
| L1M5 | 0.53% | -0.02(0.04) | 6.20E-01 | -0.06(0.48) | 1.25E-01 |
| L1ME1 | 0.49% | 0.04(0.05) | 3.96E-01 | 1.16(0.40) | 9.73E-01 |
| L1PA3 | 0.93% | 0.04(0.05) | 4.93E-01 | -0.04(0.16) | 3.57E-08 |
| L1PA4 | 0.94% | -0.05(0.05) | 3.43E-01 | 0.13(0.14) | 2.39E-04 |
| L1PA5 | 0.75% | 0.11(0.05) | 2.42E-02 | 0.49(0.16) | 7.48E-03 |
| L1PA6 | 0.45% | 0.10(0.06) | 1.17E-01 | 0.68(0.35) | 2.02E-01 |
| L1PA7 | 0.71% | 0.10(0.05) | 6.35E-02 | 0.92(0.20) | 3.50E-01 |
| L1PB1 | 0.44% | 0.09(0.03) | 1.32E-02 | 0.48(0.21) | 1.16E-01 |
| L2 | 0.49% | -0.07(0.04) | 8.20E-02 | -0.01(0.44) | 9.73E-02 |
| L2-family | 3.41% | -0.01(0.04) | 7.53E-01 | 0.81(0.17) | 5.37E-01 |
| L2a | 1.42% | 0.02(0.04) | 6.90E-01 | 0.89(0.28) | 2.71E-01 |
| L2b | 0.62% | 0.03(0.04) | 4.62E-01 | 1.30(0.47) | 8.49E-01 |
| L2c | 0.88% | -0.01(0.04) | 8.08E-01 | 0.54(0.35) | 9.28E-02 |
| MIR | 0.89% | 0.02(0.05) | 7.32E-01 | 1.01(0.46) | 9.57E-01 |
| MIRb | 1.10% | -0.01(0.04) | 7.51E-01 | 0.96(0.37) | 4.98E-01 |
| MIRc | 0.41% | 0.03(0.04) | 3.82E-01 | 1.92(0.56) | 1.91E-01 |
| TcMar-Tigger | 1.13% | 0.15(0.04) | 1.52E-04 | 1.37(0.26) | 6.17E-01 |

**Supplementary Table 10. S-LDSC results for 35 TE families/subfamilies spanning at least 0.4% of common SNPs.** We applied S-LDSC to 35 TE that capture at least 0.4% of common SNPs. These results are obtained by meta-analyzing 41 independent traits and disease conditional on baselineLD model.

| **TE** | **%SNP** | $\tau*(se)$ | $\tau*$ **(P)** | **Enrichment (se)** | **Enrichment (P)** |
|---|---|---|---|---|---|
| L1 | 17.56% | 0.16 (0.04) | 3.23E-04 | 0.62 (0.05) | 2.18E-10 |
| L1PA3 | 0.93% | 0.04 (0.05) | 4.93E-01 | -0.04 (0.16) | 3.57E-08 |
| ERV1 | 3.35% | 0.16 (0.03) | 2.11E-06 | 0.37 (0.09) | 9.08E-07 |
| L1PA4 | 0.94% | -0.05 (0.05) | 3.43E-01 | 0.13 (0.14) | 2.39E-04 |

**Supplementary Table 11. List of TE families/subfamilies spanning at least 0.4% of common SNPs with significant enrichment conditional on baseline-LD model.** We applied S-LDSC to 35 families/subfamilies spanning at least 0.4% of SNPs, to compute the enrichment and $\tau*$ of each TE meta-analyzed over 41 independent traits (47 traits) in a joint model that consist of TE and baseline-LD model (TE+baseline-LD). We reported the set of TE that have significant enrichment conditional on baseline-LD model after correcting for the number of multiple hypothesis tests ($P < \frac{0.05}{35}$).

See attached Excel file

**Supplementary Table 12. List of all 854 TE classes/families/subfamilies spanning at least 0.001% of common SNPs obtained from Repeat Masker software.** It is worth mentioning that we have 853 TE and one ALLTE. ALLTE is the combination of all TE. Thus, we have 854 rows in the Excel file.

See attached Excel file

**Supplementary Table 13. S-LDSC results for 814 TE families/subfamilies spanning less than 0.4% of common SNPs.** We computed the Expected (baseline-LD) 814 TE that capture at least 0.001% of common SNPs and less than 0.4%. These results are obtained by meta-analyzing 41 independent traits and disease. We fit the baselineLD model for each trait. Then, using S-LDSC model we can estimate the per SNP heritability (variance of each SNP). We compute the heritability of each TE by summing over the per SNP heritability of all SNPs in each corresponding TE. The miliDiv column is obtained from Repeat masker software (see URLs). We use the miliDiv as an approximation of the age of the repeat.

| TE | Enrichment (se) | Enrichment (P) |
|---|---|---|
| AmnSINE1 | 5.44 (0.32) | 1.40E-64 |
| Charlie13a | 1.36 (0.07) | 2.12E-05 |
| Charlie15a | 1.23 (0.04) | 1.01E-07 |
| Charlie19a | 1.44 (0.04) | 9.10E-14 |
| Charlie20a | 1.27 (0.06) | 4.79E-06 |
| Charlie26a | 1.43 (0.05) | 7.53E-08 |
| CR1-Hg19 | 1.33 (0.05) | 2.64E-32 |
| CR1-Mam | 1.97 (0.11) | 1.48E-31 |
| L3 | 1.11 (0.04) | 2.79E-08 |
| L3b | 1.81 (0.07) | 5.33E-41 |
| L5 | 1.71 (0.07) | 3.11E-17 |
| LFSINE-Vert | 5.54 (0.39) | 3.05E-72 |
| LTR10B1 | 2.08 (0.34) | 4.91E-09 |
| LTR14 | 2.09 (0.26) | 1.14E-05 |
| LTR90A | 1.65 (0.08) | 1.69E-09 |
| LTR90B | 1.36 (0.12) | 5.66E-05 |
| Mam-R4 | 2.67 (0.18) | 3.78E-33 |
| MamRep1161 | 1.28 (0.05) | 5.61E-08 |
| MamRep434 | 2.33 (0.13) | 1.78E-41 |
| MamSINE1 | 2.20 (0.13) | 9.11E-32 |
| MARNA | 1.29 (0.05) | 9.27E-11 |
| MER102b | 1.19 (0.04) | 9.13E-08 |
| MER102c | 1.15 (0.04) | 2.75E-05 |
| MER117 | 1.54 (0.06) | 1.16E-29 |
| MER121 | 6.21 (0.46) | 1.47E-76 |
| MER135 | 3.60 (0.32) | 9.12E-35 |
| MER81 | 1.14 (0.03) | 4.12E-05 |
| MER91A | 1.32 (0.05) | 4.45E-13 |
| MER94B | 1.57 (0.08) | 2.64E-07 |
| MIR3 | 1.42 (0.03) | 1.99E-55 |
| MLT1K | 1.18 (0.05) | 3.66E-09 |
| MLT1M | 1.42 (0.05) | 4.01E-18 |
| Plat-L3 | 2.77 (0.15) | 1.83E-60 |
| Tigger10 | 1.30 (0.06) | 1.05E-05 |
| Tigger12 | 1.45 (0.06) | 1.16E-06 |
| Tigger12A | 1.40 (0.05) | 4.24E-08 |
| Tigger12c | 1.54 (0.06) | 1.40E-15 |
| Tigger14a | 1.70 (0.07) | 5.95E-20 |
| Tigger15a | 1.44 (0.04) | 3.98E-27 |
| Tigger16b | 1.55 (0.08) | 5.06E-12 |
| UCON29 | 5.58 (0.42) | 4.49E-51 |
| X3-LINE | 2.31 (0.17) | 4.61E-19 |
| X6B-LINE | 4.93 (0.38) | 3.66E-43 |
| X7A-LINE | 1.24 (0.04) | 1.44E-06 |
| X7C-LINE | 1.83 (0.07) | 2.33E-17 |
| X8-LINE | 3.57 (0.28) | 2.75E-30 |

**Supplementary Table 14. Set of TE families/subfamilies that are enriched for the Expected (baseline-LD) heritability of disease and complex traits.**

See attached Excel file

**Supplementary Table 15. Excess overlap of 854 TE classes/families/subfamilies and functional annotations.** We compute the excess overlap between all the 853 TE classes/families/subfamilies and the baseline-LD functional annotations. We defined the excess overlap as the proportion of observed overlap between two annotations divide by the expected overlap between two annotations. Let $A$ and $B$ indicate two annotations and $|.|$ indicate the number of non-zero SNPs and we assume $M$ is the total number of common SNPs. We defined the excess overlap as follows: Excess(A,B) $= \frac{\frac{|A \cap B|}{M}}{\frac{|A|}{M}\frac{|B|}{M}}$. We compute the standard error over our estimates using block jackknife with 200 blocks (see Methods).

See attached Excel file

**Supplementary Table 16. Correlation of 854 TE classes/families/subfamilies with baseline-LD model annotations.** We compute the correlation overlap between all the 853 TE classes/families/subfamilies and the baseline-LD functional annotations. We compute the standard error over our estimates using block jackknife with 200 blocks (see Methods).

| **Trait Group** | **Trait** | Source | $N$ |
|---|---|---|---|
| Blood | Platelet Count | UKBiobank | 444,382 |
| | Red Blood Cell Count | UKBiobank | 445,174 |
| | Red Blood Cell Distribution Width | UKBiobank | 442,700 |
| | Eosinophil Count | UKBiobank | 439,938 |
| | White Blood Cell Count | UKBiobank | 444,502 |
| Autoimmune | Crohn's Disease | Jostins et al., 2012 Nature | 20,883 |
| | Rheumatoid Arthritis | Okada et al., 2014 Nature | 37,681 |
| | Ulcerative Colitis | Jostins et al., 2012 Nature | 27,432 |
| | Celiac | Dubois et al., 2010 Nat Genet | 15,283 |
| | Lupus | Bentham et al., 2015 Nat Genet | 14,267 |
| | Auto Immune Traits (Sure) | UKBiobank | 459,324 |
| Brain | Age at Menarche | UKBiobank | 242,278 |
| | BMI | Speliotes et al., 2010 Nat Genet | 122,033 |
| | BMI | UKBiobank | 457,824 |
| | Bipolar Disorder | BIP Working Group, 2011 Nat Genet | 16,731 |
| | Depressive symptoms | Okbay et al., 2016 Nat Genet | 161,460 |
| | Neuroticism | UKBiobank | 372,066 |
| | Schizophrenia | SCZ Working Group, 2014 Nature | 70,100 |
| | Smoking Status | TAG Consortium, 2010 Nat Genet | 74,035 |
| | Smoking Status | UKBiobank | 457,683 |
| | Year of education | UKBiobank | 328,917 |
| | Year of education | Rietveld et al., 2013 Science | 126,559 |

**Supplementary Table 17. List of blood, autoimmune and brain-related diseases and complex traits.** We analyzed 5 blood traits, 6 autoimmune diseases, and 8 brain-related traits. For some traits, we analyzed more than one data set.

| **Trait(T)** | **TE** | **Enr. (se)** Trait T | **$\tau *$ T (se)** | **Enr. other (se)** | **$\tau *$ other (se)** | **Expected Enr.** Trait T (se) | **Expected Enr.** other traits |
|---|---|---|---|---|---|---|---|
| Blood | ALLTE | 0.80(0.11) | 0.37(0.14) | 0.68(0.03) | 0.28(0.05) | 0.85 (0.02) | 0.84 (0.01) |
| Blood | LINE | 0.75(0.13) | 0.27(0.09) | 0.72(0.06) | 0.16(0.05) | 0.55 (0.03) | 0.57 (0.02) |
| Blood | SINE | 2.05(0.30) | 0.88(0.14) | 0.96(0.09) | 0.33(0.06) | 1.05 (0.04) | 0.93 (0.01) |
| Blood | LTR | 0.35(0.16) | 0.24(0.13) | 0.38(0.08) | 0.13(0.04) | 0.44 (0.07) | 0.53 (0.02) |
| Blood | DNA | 0.80(0.41) | 0.10(0.10) | 1.26(0.23) | 0.14(0.05) | 0.80 (0.03) | 0.87 (0.01) |
| Autoimmune | ALLTE | 0.96(0.17) | 0.67(0.29) | 0.68(0.03) | 0.28(0.05) | 0.85 (0.02) | 0.84 (0.01) |
| Autoimmune | LINE | 1.28(0.36) | 0.70(0.30) | 0.72(0.06) | 0.16(0.05) | 0.50 (0.07) | 0.57 (0.01) |
| Autoimmune | SINE | 1.33(0.69) | 0.47(0.42) | 0.96(0.09) | 0.33(0.06) | 1.09 (0.04) | 0.93 (0.01) |
| Autoimmune | LTR | 0.77(0.80) | 0.44(0.40) | 0.38(0.08) | 0.13(0.04) | 0.56 (0.04) | 0.52 (0.02) |
| Autoimmune | DNA | 2.97(2.38) | 0.60(0.55) | 1.26(0.23) | 0.14(0.05) | 0.81 (0.05) | 0.86 (0.01) |
| Brain | ALLTE | 0.64(0.04) | 0.21(0.06) | 0.69(0.03) | 0.29(0.05) | 0.84 (0.01) | 0.84 (0.01) |
| Brain | LINE | 0.63(0.10) | 0.09(0.07) | 0.71(0.06) | 0.16(0.05) | 0.64 (0.02) | 0.55 (0.02) |
| Brain | SINE | 0.86(0.11) | 0.25(0.07) | 1.00(0.09) | 0.35(0.06) | 0.87 (0.02) | 0.96 (0.01) |
| Brain | LTR | 0.73(0.14) | 0.24(0.06) | 0.39(0.09) | 0.14(0.04) | 0.42 (0.02) | 0.55 (0.02) |
| Brain | DNA | 1.32(0.28) | 0.16(0.07) | 1.29(0.23) | 0.15(0.05) | 0.90 (0.01) | 0.85 (0.01) |

**Supplementary Table 18. Trait class-specific S-LDSC results for each TE class.** In the case of blood and autoimmune traits the expected enrichment is computed using baseline-LD and blood chromatin annotation (Expected (baeline-LD+blood chromatin)) and for brain-related traits the expected enrichment is computed using baseline-LD and brain chromatin annotation (Expected (baeline-LD+brain chromatin)). Heritability and $\tau *$ are meta-analyzed over subset of traits.

| TE | Excess Blood (se) | Excess Other (se) |
|---|---|---|
| ALLTE | 0.70 (0.010) | 0.67 (0.010) |
| LINE | 0.48 (0.005) | 0.49 (0.004) |
| SINE | 0.90 (0.011) | 0.72 (0.007) |
| LTR | 0.75 (0.011) | 0.90 (0.008) |
| DNA | 0.90 (0.010) | 0.94 (0.008) |

**Supplementary Table 19. Excess overlap of each TE class with blood chromatin data and other chromatin data.** Excess overlap is computed by computing the overlap of SINE and blood active chromatin mark and non-blood active chromatin mark. We obtained the blood active chromatin regions by combining 27 blood cells and 6 chromatin mark (H3K27ac, H3K4me3, DNase, DNase-H3K27ac, DNase-H3K4me3) obtained from ChromImpute[43] applied on RoadMap[20] and we obtained the non-blood active chromatin regions by combining 100 non-blood cells and 6 chromatin mark.

| **TE** | **Enrichment Blood (se)** | **Enrichment other (se)** |
|---|---|---|
| ALU | 2.01 (0.25) | 0.97 (0.11) |
| AluJb | 1.22 (1.12) | 0.65 (0.37) |
| AluJo | 2.60 (1.44) | 1.12 (0.62) |
| AluJr | 2.11 (1.28) | 0.92 (0.75) |
| AluSc | 1.39 (1.67) | 0.47 (0.57) |
| AluSg | 1.70 (1.29) | -0.80 (0.64) |
| AluSp | 4.25 (2.06) | 1.20 (0.58) |
| AluSq2 | 0.28 (1.28) | 0.65 (0.64) |
| AluSx1 | 1.19 (0.83) | 0.89 (0.33) |
| AluSx | 1.34 (1.37) | 0.74 (0.35) |
| AluSz6 | -2.55 (2.19) | 0.66 (0.67) |
| AluSz | 1.19 (0.90) | 0.94 (0.37) |
| AluY | 0.15 (0.59) | 0.17 (0.34) |
| DNA | 0.89 (0.41) | 1.26 (0.23) |
| ERV1 | 0.16 (0.22) | 0.42 (0.10) |
| ERVL | 0.62 (0.22) | 0.59 (0.13) |
| ERVL-MaLR | 0.03 (0.28) | 0.51 (0.18) |
| hAT-Charlie | 0.46 (0.64) | 1.37 (0.38) |
| L1 | 0.76 (0.10) | 0.61 (0.06) |
| L1M5 | -0.90 (1.27) | 0.35 (0.49) |
| L1ME1 | 0.98 (0.83) | 1.30 (0.48) |
| L1PA3 | 0.07 (0.32) | -0.04 (0.18) |
| L1PA4 | 0.49 (0.30) | 0.07 (0.17) |
| L1PA5 | -0.38 (0.33) | 0.74 (0.19) |
| L1PA6 | 1.02 (0.72) | 0.59 (0.40) |
| L1PA7 | 0.95 (0.48) | 0.89 (0.22) |
| L1PB1 | 0.63 (0.48) | 0.46 (0.24) |
| L2a | 0.69 (0.67) | 0.95 (0.31) |
| L2b | -0.03 (1.80) | 1.70 (0.44) |
| L2-family | 1.19 (1.00) | -0.39 (0.49) |
| L2c | -0.57 (0.87) | 0.77 (0.39) |
| L2 | 0.61 (0.53) | 0.86 (0.18) |
| LINE | 0.81 (0.12) | 0.72 (0.06) |
| LTR | 0.39 (0.17) | 0.38 (0.08) |
| MIRb | 1.95 (1.03) | 0.86 (0.40) |
| MIR | 2.79 (1.21) | 0.79 (0.49) |
| MIRc | 0.94 (1.38) | 2.13 (0.61) |
| ALLTE | 0.83 (0.09) | 0.68 (0.03) |
| SINE | 1.96 (0.24) | 0.96 (0.09) |
| TcMar-Tigger | 1.67 (0.72) | 1.30 (0.29) |

**Supplementary Table 20. Comparison of Observed heritability enrichment of 35 TE family/subfamilies spanning at least 0.4% of common SNPs for blood traits and other traits.**

| TE | Enrichment Autoimmune (se) | Enrichment other (se) |
|---|---|---|
| ALU | 2.01 (0.25) | 0.97 (0.11) |
| AluJb | 1.22 (1.12) | 0.65 (0.37) |
| AluJo | 2.60 (1.44) | 1.12 (0.62) |
| AluJr | 2.11 (1.28) | 0.92 (0.75) |
| AluSc | 1.39 (1.67) | 0.47 (0.57) |
| AluSg | 1.70 (1.29) | -0.80 (0.64) |
| AluSp | 4.25 (2.06) | 1.20 (0.58) |
| AluSq2 | 0.28 (1.28) | 0.65 (0.64) |
| AluSx1 | 1.19 (0.83) | 0.89 (0.33) |
| AluSx | 1.34 (1.37) | 0.74 (0.35) |
| AluSz6 | -2.55 (2.19) | 0.66 (0.67) |
| AluSz | 1.19 (0.90) | 0.94 (0.37) |
| AluY | 0.15 (0.59) | 0.17 (0.34) |
| DNA | 0.89 (0.41) | 1.26 (0.23) |
| ERV1 | 0.16 (0.22) | 0.42 (0.10) |
| ERVL | 0.62 (0.22) | 0.59 (0.13) |
| ERVL-MaLR | 0.03 (0.28) | 0.51 (0.18) |
| hAT-Charlie | 0.46 (0.64) | 1.37 (0.38) |
| L1 | 0.76 (0.10) | 0.61 (0.06) |
| L1M5 | -0.90 (1.27) | 0.35 (0.49) |
| L1ME1 | 0.98 (0.83) | 1.30 (0.48) |
| L1PA3 | 0.07 (0.32) | -0.04 (0.18) |
| L1PA4 | 0.49 (0.30) | 0.07 (0.17) |
| L1PA5 | -0.38 (0.33) | 0.74 (0.19) |
| L1PA6 | 1.02 (0.72) | 0.59 (0.40) |
| L1PA7 | 0.95 (0.48) | 0.89 (0.22) |
| L1PB1 | 0.63 (0.48) | 0.46 (0.24) |
| L2a | 0.69 (0.67) | 0.95 (0.31) |
| L2b | -0.03 (1.80) | 1.70 (0.44) |
| L2-family | 1.19 (1.00) | -0.39 (0.49) |
| L2c | -0.57 (0.87) | 0.77 (0.39) |
| L2 | 0.61 (0.53) | 0.86 (0.18) |
| LINE | 0.81 (0.12) | 0.72 (0.06) |
| LTR | 0.39 (0.17) | 0.38 (0.08) |
| MIRb | 1.95 (1.03) | 0.86 (0.40) |
| MIR | 2.79 (1.21) | 0.79 (0.49) |
| MIRc | 0.94 (1.38) | 2.13 (0.61) |
| ALLTE | 0.83 (0.09) | 0.68 (0.03) |
| SINE | 1.96 (0.24) | 0.96 (0.09) |
| TcMar-Tigger | 1.67 (0.72) | 1.30 (0.29) |

**Supplementary Table 21. Comparison of Observed heritability enrichment of 35 TE families/subfamilies spanning at least 0.4% of common SNPs for autoimmune traits and other traits**

| TE | Enrichment Brain traits(se) | Enrichment other traits(se) |
|---|---|---|
| ALU | 0.82 (0.15) | 1.00 (0.10) |
| AluJb | 0.97 (0.55) | 0.71 (0.37) |
| AluJo | 1.83 (0.86) | 1.12 (0.62) |
| AluJr | 1.77 (0.84) | 0.92 (0.74) |
| AluSc | 1.63 (0.88) | 0.44 (0.57) |
| AluSg | 0.03 (1.04) | -0.76 (0.64) |
| AluSp | 0.06 (0.75) | 1.28 (0.58) |
| AluSq2 | -1.00 (0.76) | 0.71 (0.64) |
| AluSx1 | 0.48 (0.50) | 0.96 (0.33) |
| AluSx | 1.12 (0.50) | 0.76 (0.36) |
| AluSz6 | 0.77 (0.90) | 0.83 (0.59) |
| AluSz | 0.27 (0.66) | 0.96 (0.37) |
| AluY | -1.07 (0.35) | 0.19 (0.34) |
| DNA | 1.32 (0.28) | 1.29 (0.23) |
| ERV1 | 0.73 (0.15) | 0.45 (0.10) |
| ERVL | 0.66 (0.26) | 0.58 (0.13) |
| ERVL-MaLR | 0.32 (0.30) | 0.50 (0.18) |
| hAT-Charlie | 0.94 (0.65) | 1.39 (0.37) |
| L1 | 0.55 (0.12) | 0.60 (0.06) |
| L1M5 | 1.05 (1.02) | 0.32 (0.48) |
| L1ME1 | 0.48 (0.98) | 1.26 (0.44) |
| L1PA3 | -0.01 (0.30) | -0.04 (0.18) |
| L1PA4 | -0.40 (0.33) | 0.06 (0.17) |
| L1PA5 | 0.73 (0.28) | 0.76 (0.19) |
| L1PA6 | -0.57 (0.68) | 0.59 (0.40) |
| L1PA7 | 0.90 (0.34) | 0.91 (0.22) |
| L1PB1 | 0.19 (0.40) | 0.48 (0.24) |
| L2a | 1.08 (0.64) | 0.93 (0.31) |
| L2b | 2.01 (0.73) | 1.70 (0.44) |
| L2 | -0.62 (0.81) | -0.37 (0.48) |
| L2c | 0.97 (0.71) | 0.78 (0.39) |
| L2-family | 0.93 (0.41) | 0.85 (0.18) |
| LINE | 0.63 (0.10) | 0.71 (0.06) |
| LTR | 0.73 (0.14) | 0.39 (0.09) |
| MIRb | 2.04 (0.70) | 1.12 (0.47) |
| MIR | 1.76 (0.62) | 0.76 (0.49) |
| MIRc | 3.57 (0.98) | 2.11 (0.61) |
| RepeatMask | 0.64 (0.04) | 0.69 (0.03) |
| SINE | 0.86 (0.11) | 1.00 (0.09) |
| TcMar-Tigger | 2.12 (0.45) | 1.33 (0.29) |

**Supplementary Table 22. Comparison of Observed heritability enrichment of 35 TE families/subfamilies spanning at least 0.4% of common SNPs for brain traits and other traits.**

| TE | Repeat Class | Expected Enr. Blood traits(se) | Expected Enr other traits(se) | Enr P Blood vs. other traits |
|---|---|---|---|---|
| FLAM-C | SINE | 1.49 (0.11) | 1.02 (0.04) | 1.43E-05 |
| FRAM | SINE | 1.38 (0.09) | 0.94 (0.04) | 6.71E-06 |
| Harlequin-int | LTR | 0.82 (0.09) | 0.35 (0.07) | 2.48E-05 |
| HERV15-int | LTR | 2.52 (0.34) | 0.73 (0.15) | 8.33E-07 |
| HERV3-int | LTR | 1.58 (0.19) | 0.50 (0.14) | 1.49E-06 |
| HERVFH21-int | LTR | 1.25 (0.28) | -0.01 (0.13) | 2.56E-05 |
| LTR13A | LTR | 2.50 (0.26) | 1.34 (0.13) | 4.05E-05 |
| LTR14 | LTR | 2.88 (0.34) | 1.18 (0.16) | 3.98E-06 |
| LTR15 | LTR | 2.94 (0.33) | 0.97 (0.12) | 7.53E-09 |
| LTR19C | LTR | 2.15 (0.22) | 0.86 (0.13) | 2.03E-07 |
| LTR21A | LTR | 2.26 (0.27) | 0.76 (0.15) | 7.82E-07 |
| LTR24B | LTR | 1.44 (0.13) | 0.67 (0.09) | 4.34E-07 |
| LTR2B | LTR | 2.58 (0.22) | 1.10 (0.15) | 1.48E-08 |
| LTR2C | LTR | 1.84 (0.22) | 0.85 (0.10) | 1.87E-05 |
| LTR46 | LTR | 1.60 (0.16) | 0.53 (0.10) | 5.61E-09 |
| LTR4 | LTR | 3.98 (0.47) | 1.57 (0.27) | 3.99E-06 |
| LTR54 | LTR | 1.25 (0.09) | 0.76 (0.05) | 2.06E-06 |
| LTR61 | LTR | 1.53 (0.25) | 0.39 (0.12) | 2.04E-05 |
| LTR62 | LTR | 1.54 (0.17) | 0.61 (0.10) | 1.03E-06 |
| LTR65 | LTR | 1.48 (0.11) | 0.91 (0.04) | 8.95E-07 |
| LTR75-1 | LTR | 1.30 (0.18) | 0.48 (0.06) | 7.68E-06 |
| LTR7C | LTR | 1.45 (0.13) | 0.69 (0.12) | 5.30E-06 |
| MER34C2 | LTR | 1.37 (0.12) | 0.81 (0.05) | 1.26E-05 |
| MER51B | LTR | 1.06 (0.08) | 0.60 (0.07) | 3.95E-06 |
| MER51D | LTR | 1.53 (0.17) | 0.39 (0.10) | 5.54E-09 |
| MER57E3 | LTR | 5.19 (0.47) | 1.00 (0.26) | 4.75E-15 |
| MER65C | LTR | 0.98 (0.09) | 0.56 (0.06) | 2.04E-05 |

**Supplementary Table 23. Comparison of Expected (baseline-LD+blood chromatin) enrichment of TE families/subfamilies in blood cell-type traits and non-blood cell-type traits.** These are TE families/subfamilies that are significantly different enriched between blood and non-blood traits.

| TE | Repeat Class | Expected Enr. Autoimmune traits(se) | Expected Enr other traits(se) | Enr P Autoimmune vs. other traits |
|---|---|---|---|---|
| FLAM-C | SINE | 1.54 (0.11) | 1.04 (0.04) | 1.38E-05 |
| FRAM | SINE | 1.28 (0.06) | 0.97 (0.04) | 1.83E-05 |
| HERVFH21-int | LTR | 4.14 (0.84) | 0.03 (0.12) | 6.53E-07 |
| LOR1a | LTR | 1.13 (0.12) | 0.52 (0.05) | 1.71E-06 |
| LTR10A | LTR | 2.77 (0.37) | 0.90 (0.13) | 1.14E-06 |
| LTR10B | LTR | 2.80 (0.40) | 0.81 (0.14) | 1.27E-06 |
| LTR10F | LTR | 1.50 (0.23) | 0.48 (0.11) | 3.82E-05 |
| LTR12F | LTR | 1.65 (0.29) | 0.34 (0.06) | 3.49E-06 |
| LTR13A | LTR | 3.36 (0.44) | 1.38 (0.13) | 7.18E-06 |
| LTR19A | LTR | 1.67 (0.25) | 0.61 (0.10) | 3.85E-05 |
| LTR21A | LTR | 4.12 (0.76) | 0.84 (0.15) | 1.11E-05 |
| LTR2 | LTR | 1.61 (0.16) | 0.79 (0.08) | 2.66E-06 |
| LTR4 | LTR | 7.54 (1.09) | 1.62 (0.26) | 6.66E-08 |
| LTR62 | LTR | 1.71 (0.21) | 0.65 (0.10) | 3.82E-06 |
| LTR76 | LTR | 3.06 (0.49) | 1.00 (0.17) | 3.65E-05 |
| LTR7C | LTR | 1.85 (0.26) | 0.72 (0.12) | 4.06E-05 |
| MER41B | LTR | 1.70 (0.10) | 0.93 (0.08) | 1.54E-09 |
| MER41D | LTR | 1.55 (0.21) | 0.55 (0.07) | 1.91E-06 |
| MER41E | LTR | 1.96 (0.27) | 0.65 (0.08) | 1.84E-06 |
| MER51D | LTR | 2.22 (0.33) | 0.45 (0.11) | 2.09E-07 |
| MER57F | LTR | 1.88 (0.32) | 0.54 (0.06) | 2.40E-05 |
| MER83B | LTR | 2.67 (0.42) | 0.96 (0.09) | 3.82E-05 |
| MLT1F | LTR | 1.35 (0.07) | 0.76 (0.03) | 2.40E-13 |
| MLT1G3 | LTR | 1.13 (0.08) | 0.70 (0.03) | 4.27E-07 |
| MLT1J | LTR | 1.18 (0.04) | 0.82 (0.03) | 3.02E-15 |
| MLT1K | LTR | 1.19 (0.04) | 0.95 (0.03) | 1.34E-06 |
| MLT1L | LTR | 1.21 (0.06) | 0.85 (0.03) | 8.31E-08 |

**Supplementary Table 24. Comparison of Expected (baseline-LD+blood chromatin) enrichment of TE in autoimmune traits and other traits.** These are TE families/subfamilies that are significantly different enriched between autoimmune and other traits.

See attached Excel file

**Supplementary Table 25. Comparison of Expected (baseline-LD+brain chromatin) enrichment of TE in brain-related traits and other traits.** These are TE families/subfamilies that are significantly different enriched between brain-related traits and other traits.

| Window | Repeat | $\tau*(se)$ | $\tau*$ (P) | Enrichment (se) | Enrichment (P) |
|---|---|---|---|---|---|
| window-500bp | ALU | 0.49 (0.06) | 1.12E-15 | 1.18 (0.12) | 2.35E-01 |
| | DNA | 0.15 (0.04) | 8.64E-04 | 1.23 (0.19) | 6.68E-01 |
| | ERVK | 0.09 (0.04) | 1.88E-02 | -0.66 (0.26) | 3.50E-05 |
| | L1 | 0.16 (0.04) | 3.23E-04 | 0.62 (0.05) | 2.18E-10 |
| | LINE | 0.18 (0.05) | 1.73E-04 | 0.73 (0.05) | 4.66E-07 |
| | LTR | 0.14 (0.03) | 2.90E-05 | 0.38 (0.07) | 2.81E-08 |
| | ALLTE | 0.31 (0.05) | 1.14E-11 | 0.72 (0.03) | 3.68E-12 |
| | SINE | 0.43 (0.06) | 3.09E-12 | 1.18 (0.11) | 2.73E-01 |
| window-padding | ALU | 0.30 (0.09) | 5.49E-04 | 1.08 (0.14) | 8.52E-01 |
| | DNA | 0.09 (0.06) | 1.52E-01 | 1.32 (0.20) | 4.13E-01 |
| | ERVK | 0.34 (0.10) | 4.08E-04 | -0.55 (0.30) | 1.50E-03 |
| | L1 | -0.04 (0.09) | 6.94E-01 | 0.65 (0.05) | 5.56E-08 |
| | LINE | 0.03 (0.06) | 5.78E-01 | 0.70 (0.05) | 3.54E-07 |
| | LTR | 0.22 (0.06) | 2.50E-04 | 0.47 (0.08) | 2.33E-06 |
| | ALLTE | 0.17 (0.06) | 1.02E-02 | 0.69 (0.03) | 4.81E-13 |
| | SINE | 0.22 (0.08) | 5.14E-03 | 1.04 (0.12) | 8.23E-01 |

**Supplementary Table 26. Choice of window size does not impact S-LDSC heritability enrichment results.** In our main analysis, for each TE class/family we add an additional annotation that consist of a 500bp window around the TE. We refer to this annotation as window-500. To make sure our result is not biased to the window size (e.g., 500bp), we consider a model where we add four different window sizes (100, 200, 500, and 1000bp) around each TE annotation.

| Annotation | Expected (%SNPs) | $\tau*(se)$ | $\tau*$ (P) | Enrichment (se) | Enrichment (P) |
|---|---|---|---|---|---|
| ALLTE | 47.08 | 0.31 (0.04) | 6.92E-13 | 0.77 (0.02) | 3.86E-09 |
| LINE | 20.04 | 0.23 (0.05) | 4.04E-05 | 0.82 (0.05) | 1.72E-03 |
| SINE | 10.75 | 0.59 (0.06) | 1.57E-20 | 1.80 (0.14) | 1.78E-06 |
| DNA | 3.71 | 0.20 (0.04) | 4.61E-06 | 1.21 (0.15) | 7.94E-01 |
| LTR | 9.98 | 0.16 (0.04) | 1.31E-05 | 0.39 (0.07) | 8.81E-08 |

**Supplementary Table 27. S-LDSC results for main annotations restricted to SNPs with mappability of 1.** We report the heritability enrichment and $\tau*$ for four main TE classes (LINE, SINE, LTR, and DNA), and ALLTE restricted to the 85% of SNPs with mappability of 1. Results are meta-analyzed across 41 independent traits. Results were very similar to analyses using all SNPs (Figure 1 and Supplementary Table 3).

### Supplementary Figures

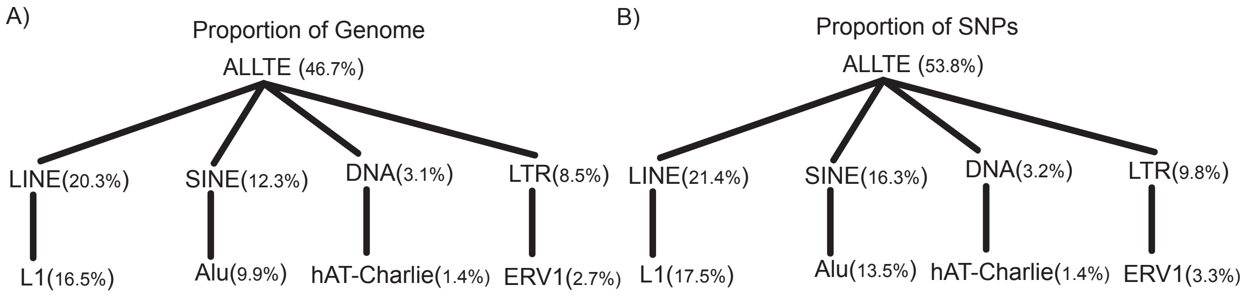


**Supplementary Figure 1. Proportion of genome and proportion of SNPs spanned by each TE class and corresponding TE families.** We have four main TE classes (LINE, SINE, DNA, and LTR). For each main TE class, we illustrated the family that captures the largest proportion of that TE class (L1 for LINE, Alu for SINE, hAT-Charlie for DNA, and ERV1 for LTR). (A) Proportion of genome that is captured by each TE. (B) Proportion of common SNPs that is captured by each TE.

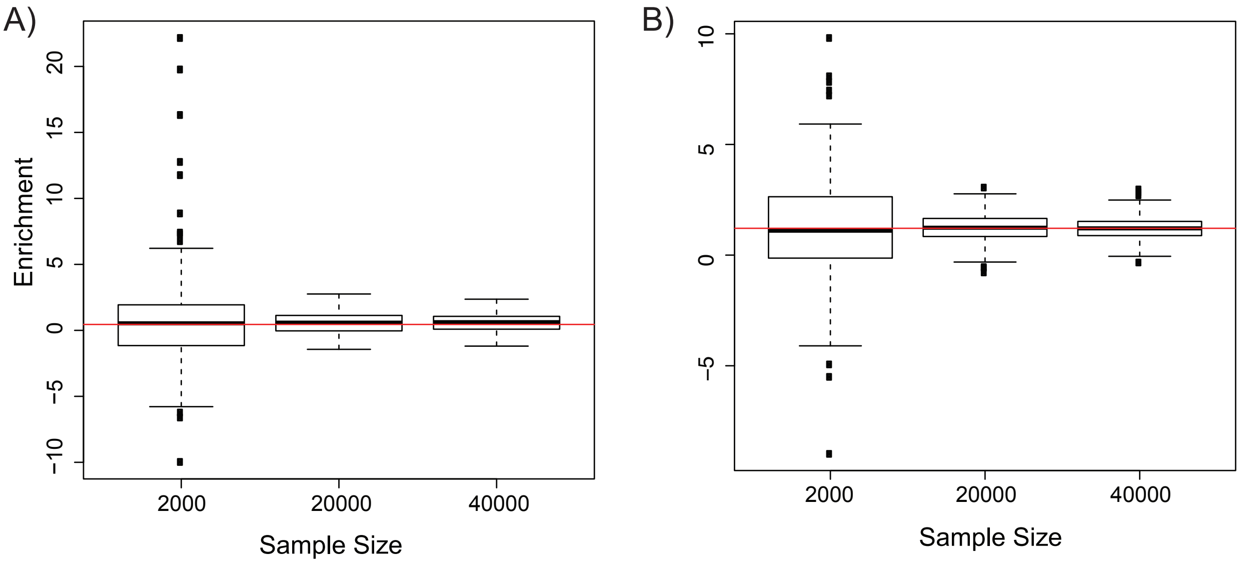


**Supplementary Figure 2. S-LDSC produces unbiased estimates of enrichment in simulations with TE annotations.** We simulated enrichment for different TE annotations. Then, we simulated the summary statistics for 2,000, 20,000, and 40,000 individuals where the genotypes are obtained from UK Biobank. After simulating the summary statistics, we applied S-LDSC conditional on baseline-LD model and the TE annotation. Regression SNPs in S-LDSC were obtained from the HapMap Project phase 3[42] (see URLs). These SNPs are well-imputed SNPs. SNPs with marginal association statistics larger than 80 or larger than 0.001N and SNPs that are in the major histocompatibility complex (MHC) region were excluded from all the analyses. Reference SNPs were obtained using the European samples in 1000G[41]. Heritability SNPs, which are used to estimate $h_g^2$, were common variants (MAF$\geq$0.05) in the set of reference SNPs. A) We simulated enrichment for LTR where we set LTR true enrichment value to be 0.44, which is close to the estimated enrichment for LTR in real dataset. B) We simulated enrichment for SINE where we set SINE true enrichment value to be 1.22, which is close to the estimated enrichment for SINE in real dataset. The red horizontal line indicates the true simulated enrichment for each TE. The X-axis is the number of individuals and the Y-axis is the estimated enrichment from S-LDSC.

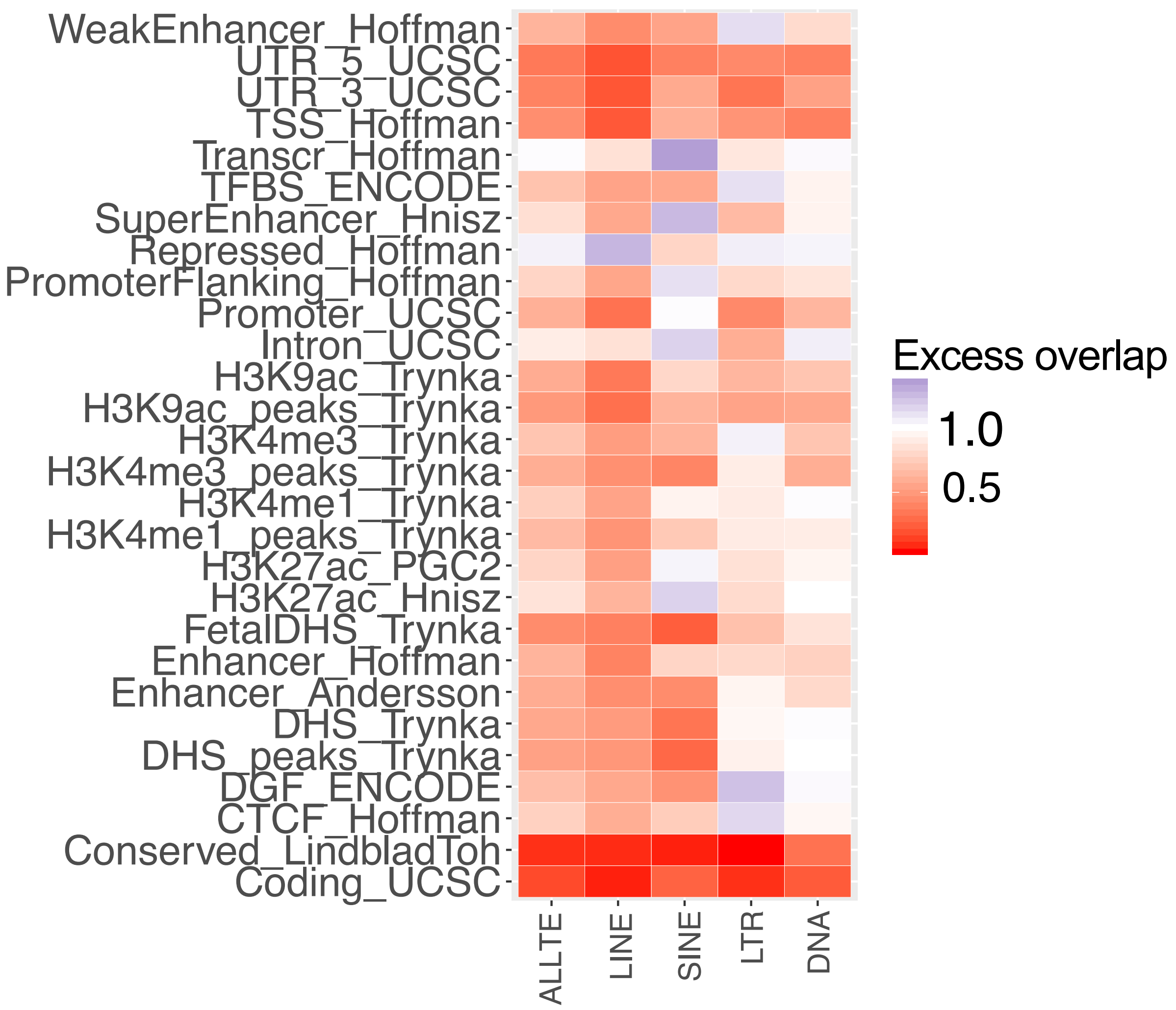


**Supplementary Figure 3. Excess overlap between each TE class and functional annotations.** The X-axis is the set of 5 TE (All, LINE, SINE, LTR, and DNA) and the Y-axis is the set of functional annotations in baseline model. We defined the excess overlap as the proportion of observed overlap between two annotations divide by the expected overlap between two annotations. Let $A$ and $B$ indicate two annotations and $|.|$ indicate the number of non-zero SNPs and we assume $M$ is the total number of common SNPs. We defined the excess overlap as follows: Excess(A,B) $= \frac{\frac{|A \cap B|}{M}}{\frac{|A|}{M}\frac{|B|}{M}}$.

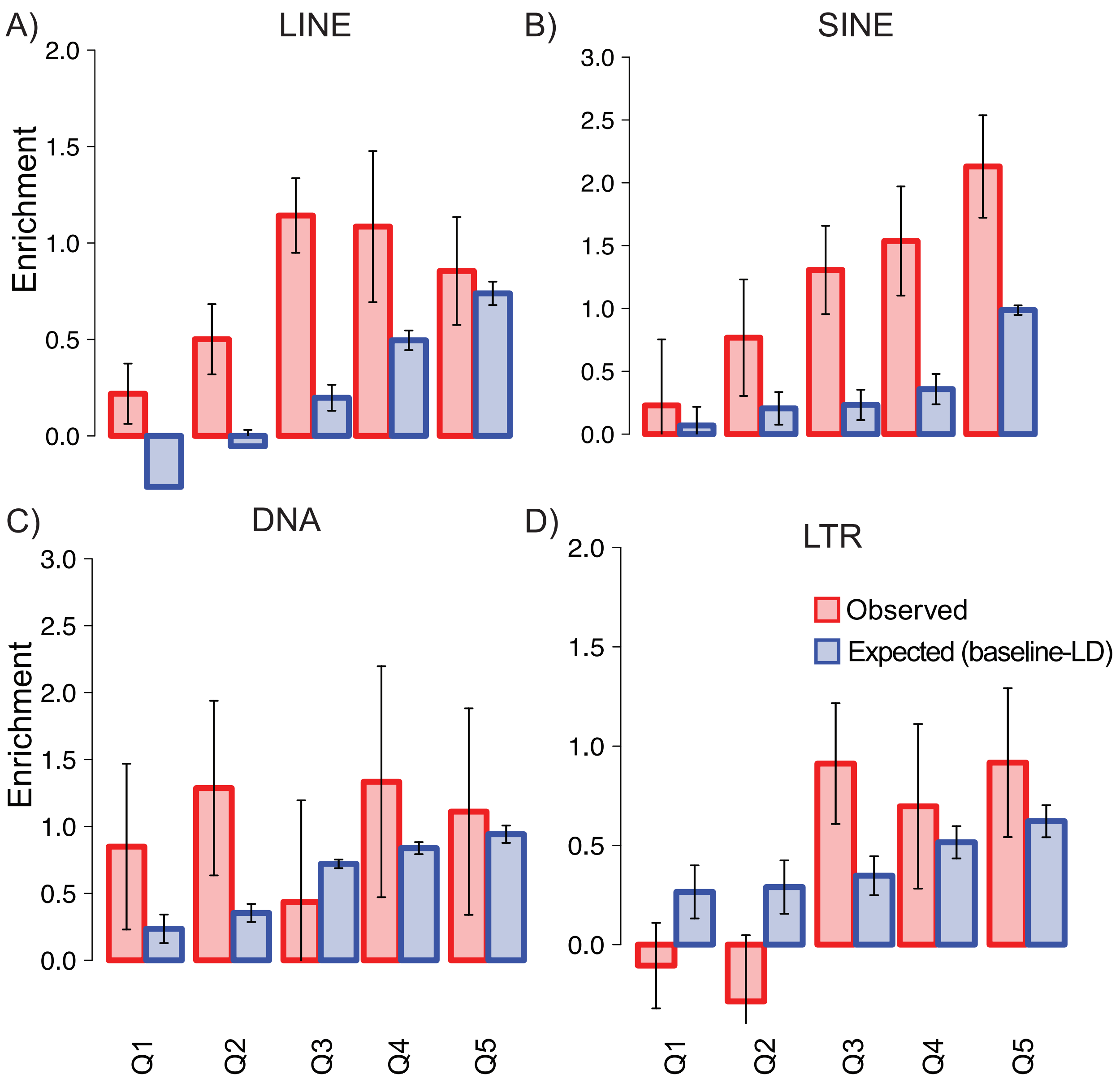


**Supplementary Figure 4. S-LDSC results for all TE in different age quintiles.** We report the Observed and Expected (baseline-LD) enrichment of SNPs in ALLTE for five different age quintiles where Q1 is the youngest and Q5 is the oldest. We computed the Observed enrichment conditional on ALLTE and the baseline-LD model. Error bars represent 95% confidence intervals.

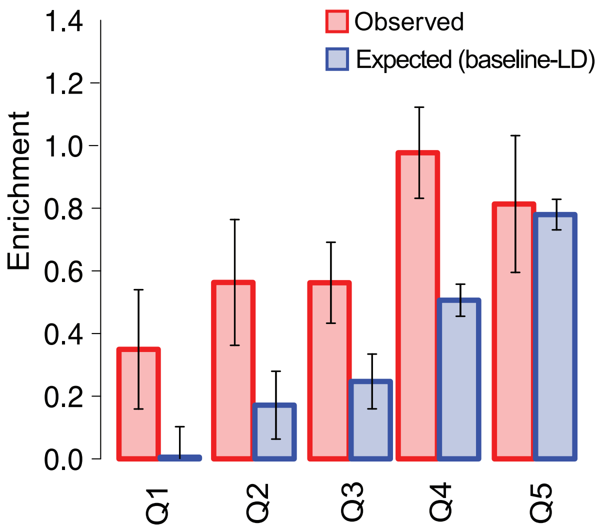


**Supplementary Figure 5. S-LDSC results for each TE class in different age quintiles.** We report the Observed and Expected (baseline-LD) enrichment of SNPs in each TE class for five different age quintiles where Q1 is the youngest and Q5 is the oldest. We computed the Observed enrichment conditional on ALLTE and the baseline-LD model. Error bars represent 95% confidence intervals. Numerical results are reported in Supplementary Table 7.

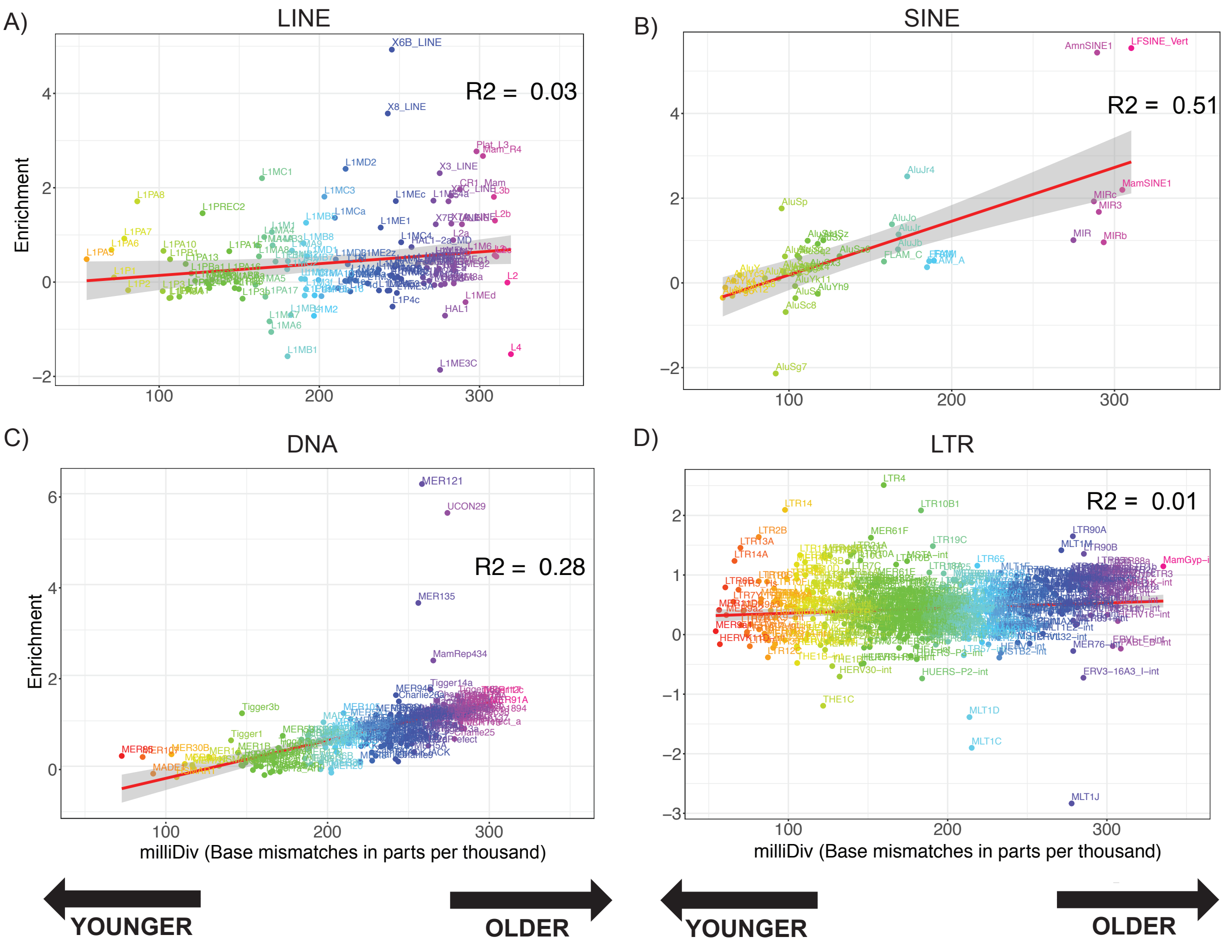


**Supplementary Figure 6. S-LDSC results for individual TE families/subfamilies in each TE class as a function of age.** Meta-analysis results of expected (baseline-LD) enrichment for (A) LINE, (B) SINE, (C) DNA, and (D) LTR across 41 traits. The X-axis is the age in miliDiv that is obtained from Repeat masker software (see URLs). We use the miliDiv as an approximation of the age of TE. The Y-axis is the expected (baseline-LD) enrichment for each TE family/subfamily. The red line indicates a linear fit between the miliDiv and enrichment of each TE family/subfamily. Numerical results are reported in Supplementary Table 8.

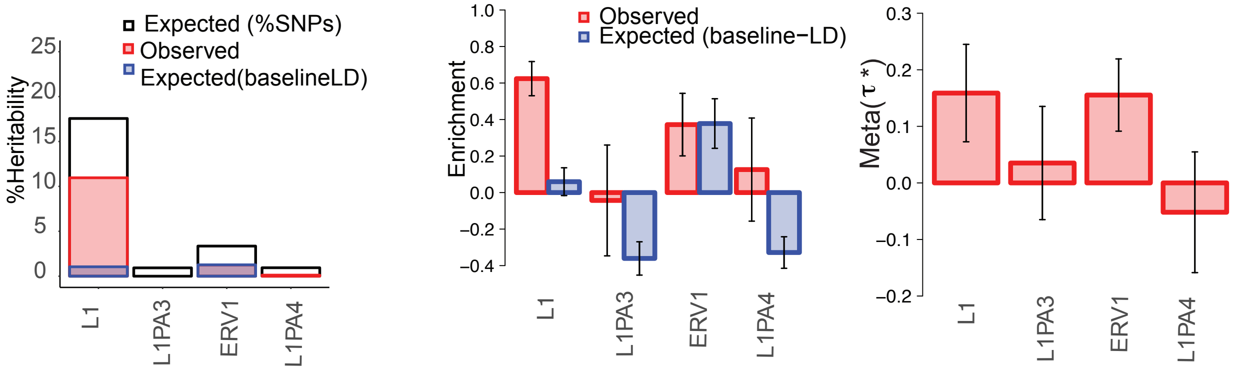


**Supplementary Figure 7. TE families/subfamilies spanning at least 0.4% of common SNPs that are significantly depleted for disease heritability.** A) Expected and observed %heritability (proportion of trait heritability) captured by each four TE families/subfamiles (L1, L1PA3, ERV1, L1PA4). Meta-analysis results across 41 traits of (B) enrichment and (C) $\tau*$ for all TE families/subfamiles that have significant enrichment conditional on the baseline-LD model. Error bars represent 95% confidence intervals. Numerical results are reported in Supplementary Table 11.

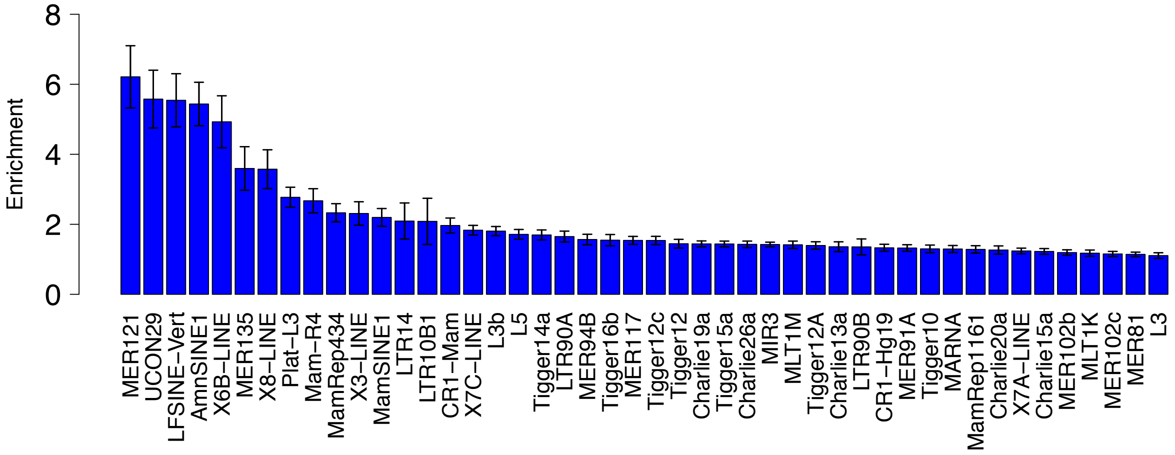


**Supplementary Figure 8. TE families/subfamilies spanning less than 0.4% of common SNPs that are significantly enriched for disease heritability.** We compute the trait enrichment of TE families/subfamilies that span less than 0.4% of SNPs by fitting a baselineLD model. Then we compute the per-SNP heritability based on $\tau$ estimated from baselineLD model. Enrichment of a TE is computed by summing over the per-SNP heritability of all SNPs that belong to TE. The X-axis is the set of TE and Y-axis is the trait enrichment. We detected 46 TE that are enriched for the heritability of disease and complex traits. Error bars represent 95% confidence intervals. Numerical results are reported in Supplementary Table 14.

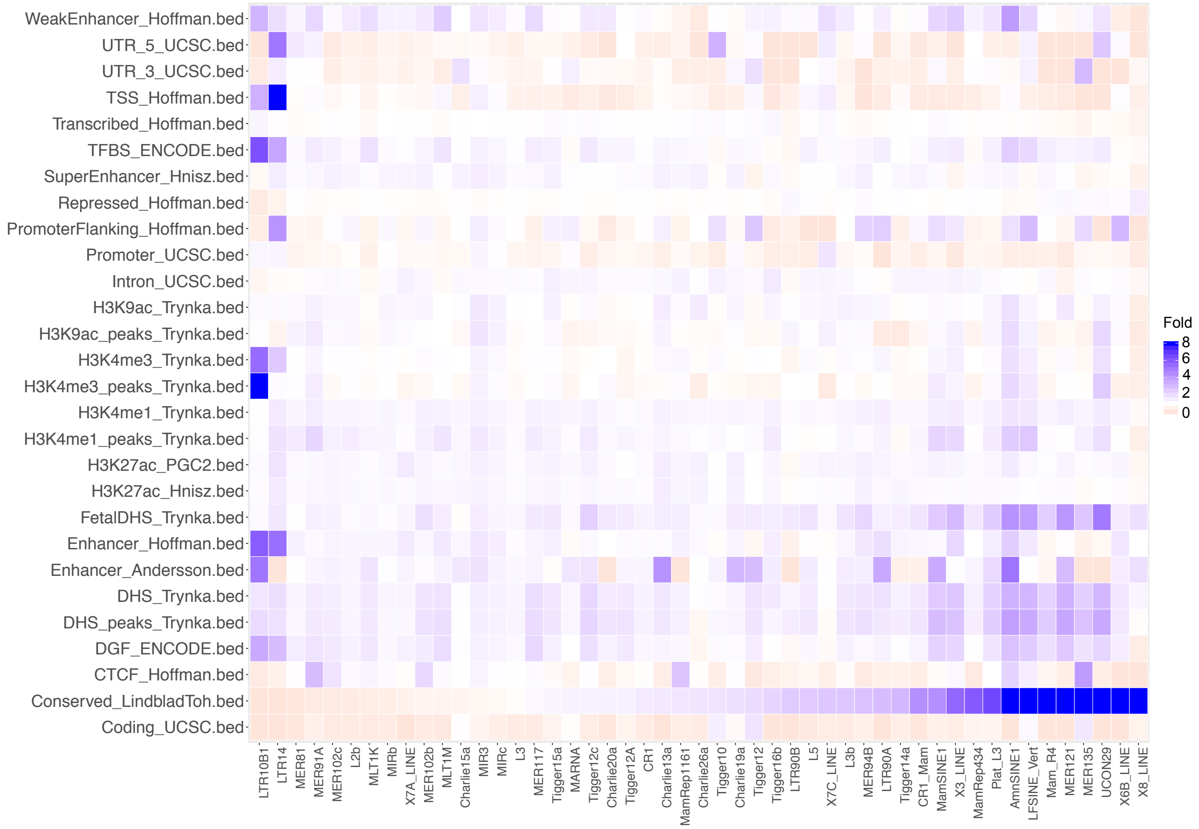


**Supplementary Figure 9. Excess overlap between TE families/subfamilies spanning less than 0.4% of common SNPs that are significantly enriched for disease heritability and functional annotations.** We only considered 46 TE that are enriched for trait heritability. The X-axis is the set of TE and the Y-axis is the set of functional annotations in baseline model. We defined the excess overlap as the proportion of observed overlap between two annotations divide by the expected overlap between two annotations. Let $A$ and $B$ indicate two annotations and $|.|$ indicate the number of non-zero SNPs and we assume $M$ is the total number of common SNPs. We defined the excess overlap as follows: Excess(A,B) = $\frac{\frac{|A \cap B|}{M}}{\frac{|A|}{M}\frac{|B|}{M}}$. We compute the standard error over our estimates using block jackknife with 200 blocks (see Methods). Numerical results are reported in Supplementary Table 15.

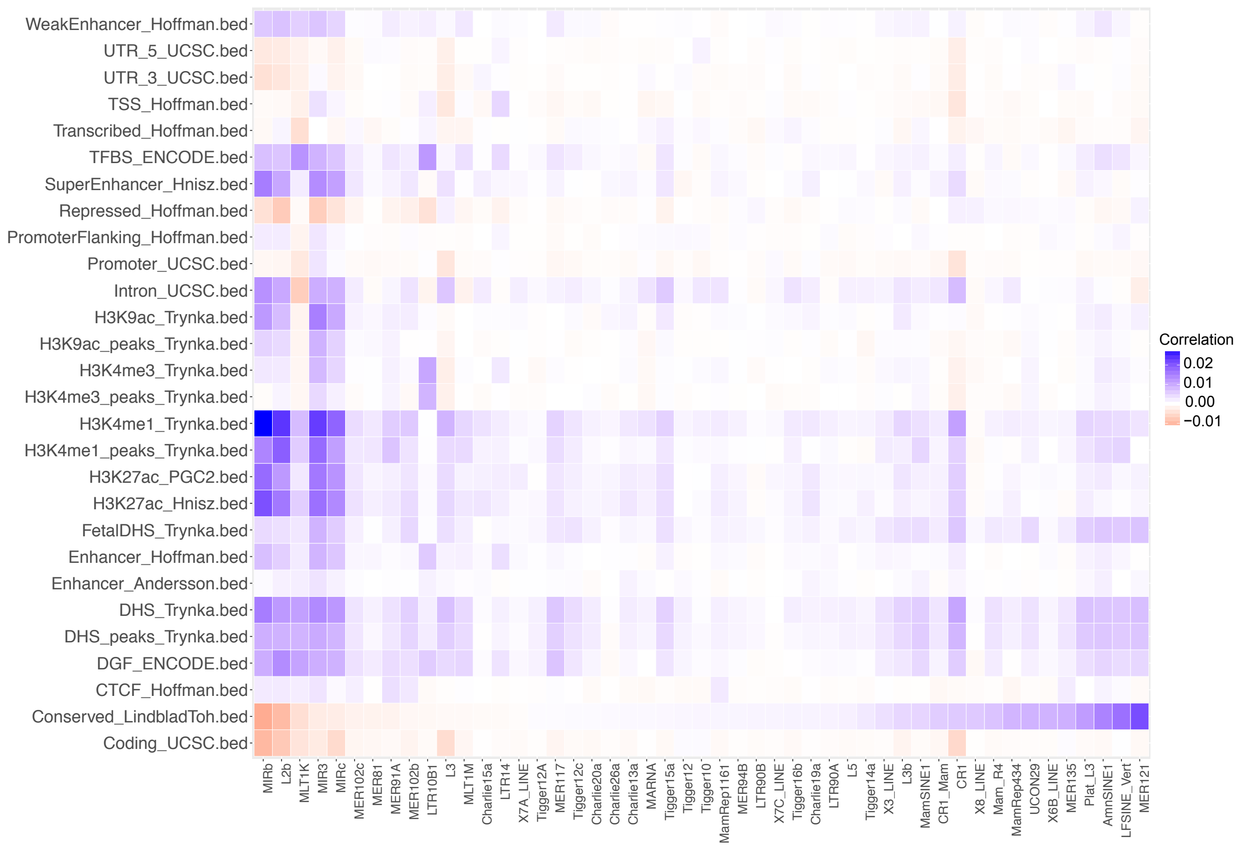


**Supplementary Figure 10. Correlation between TE families/subfamilies spanning less than 0.4% of common SNPs that are significantly enriched for disease heritability and functional annotations.** We only considered 46 TE that are enriched for trait heritability. We use correlation as a measure to compute similarity between two annotations. We compute the correlation between each TE and each baseline annotation. We compute the standard error over our estimates using block jackknife with 200 blocks (see Methods). We sort the TE in an increasing order of conserved annotation correlation with each TE. Numerical results are reported in Supplementary Table 16.

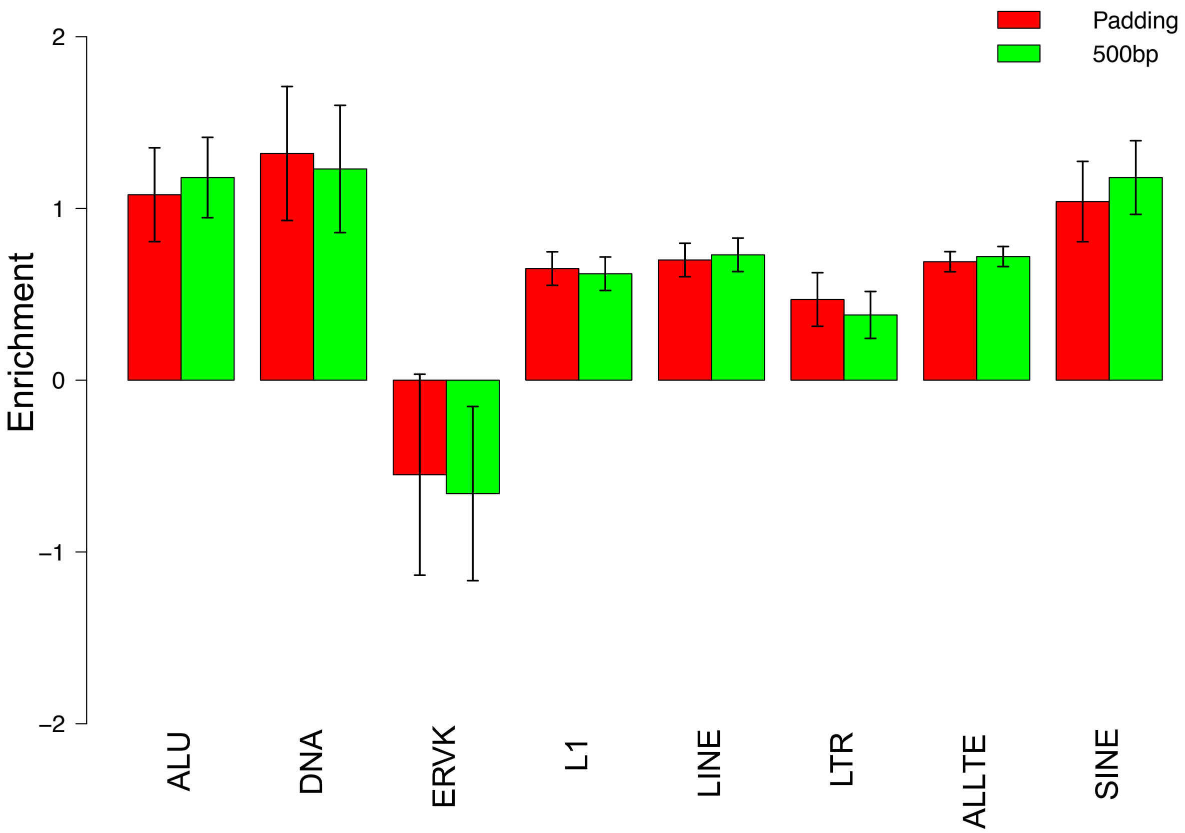


**Supplementary Figure 11. Comparing S-LDSC enrichment of TE classes/families for different window sizes.** In our main analysis, for each TE we add an additional annotation that consist of a 500bp window around the TE. We refer to this annotation as window-500. To make sure our result is not biased to the window size (e.g., 500bp), we consider a model where we add four different window sizes (100, 200, 500, and 1000bp) around each TE annotation. We refer to this result as window-padding. Numerical results are reported in Supplementary Table 26.
